# Genomic signatures of isolation, hybridization, and selection during speciation of island finches

**DOI:** 10.1101/2022.01.05.474904

**Authors:** Martin Stervander, Martim Melo, Peter Jones, Bengt Hansson

**Affiliations:** Department of Biology, Lund University, Ecology Building, 223 62 Lund, Sweden; CIBIO, Research Centre in Biodiversity and Genetic Resources, InBio, Associated Laboratory, University of Porto, Campus Agrário de Vairão, Rua Padre Armando Quintas 7, 4485-661 Vairão, Portugal; MHNC-UP, Natural History and Science Museum of the University of Porto, Praça Gomes Teixeira, 4099-002 Porto, Portugal; FitzPatrick Institute of African Ornithology, University of Cape Town, Private Bag X3, Rondebosch 7701, Cape Town, South Africa; Institute of Evolutionary Biology, University of Edinburgh, Edinburgh EH9 3JT, Scotland

**Keywords:** adaptation, bill morphology, birds, colonization, introgression, resource-driven selection

## Abstract

Sister species occurring sympatrically on islands are rare and offer unique opportunities to understand how speciation can proceed in the face of gene flow. The São Tomé grosbeak is a massive-billed, ‘giant’ finch endemic to the island of São Tomé in the Gulf of Guinea, where it has diverged from its co-occurring sister species the Príncipe seedeater, an average-sized finch that also inhabits two neighbouring islands. Here, we show that the grosbeak carries a large number of unique alleles different from all three Príncipe seedeater populations, but also shares many alleles with the sympatric São Tomé population of the seedeater, a genomic signature signifying divergence in isolation as well as subsequent introgressive hybridization. Furthermore, genomic segments that remain unique to the grosbeak are situated close to genes, including genes that determine bill morphology, suggesting the preservation of adaptive variation through natural selection during divergence with gene flow. This study reveals a complex speciation process whereby genetic drift, introgression, and selection during periods of isolation and secondary contact all have shaped the diverging genomes of these sympatric island endemic finches.

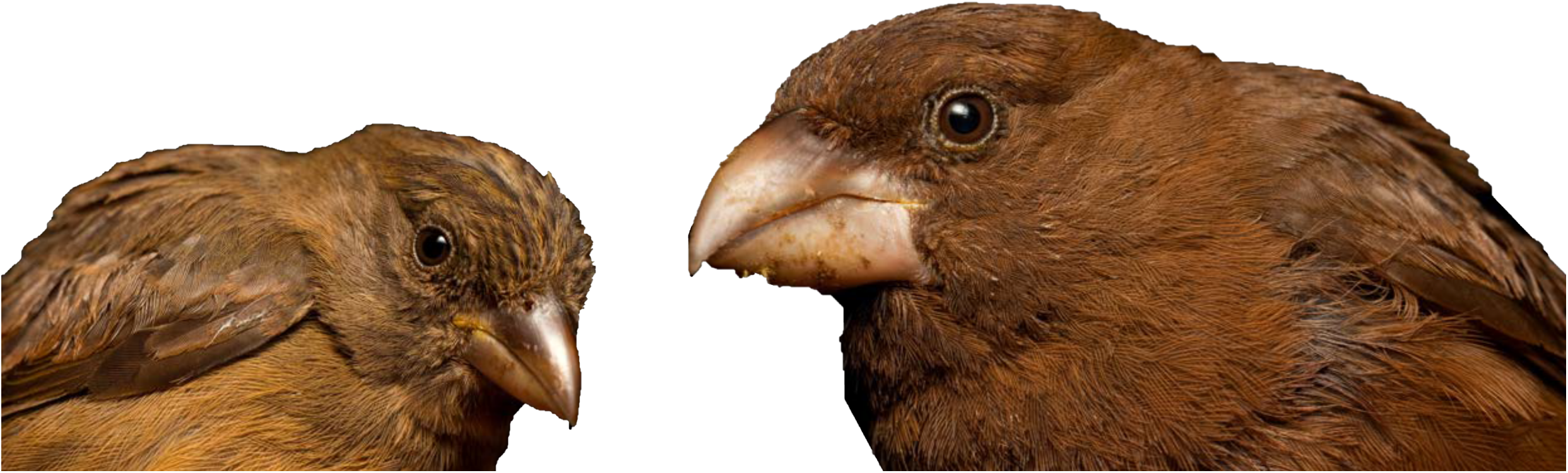

## INTRODUCTION

Adaptation to distinct ecological conditions can cause population differentiation and lead to the evolution of reproductive isolation^1-3^. Population differentiation is counteracted, however, by the homogenizing effect of gene flow^4,5^, and although ‘speciation with gene flow’ is accepted as occurring in nature^6^, divergence happens much more readily when populations experience some degree of geographical isolation^7^, as in the archipelago radiation of the Darwin’s finches^8,9^.

Understanding the process of speciation remains challenging because complex demographic scenarios and selection regimes may interact during divergence and leave similar signatures in the genome^10^. Pairs of closely related species present in small, isolated, habitat islands—like oceanic islands, caves and lakes—constitute the most tractable models to study speciation with gene flow in the wild^11^, and hence to gain insights into the role of ecology, isolation, and introgression during speciation. Sympatric sister species of island birds are rare^12^, but here, we report on the evolution of two endemic sister species of birds co-occurring on the island of São Tomé in the Gulf of Guinea, Central Africa: the enigmatic and Critically Endangered São Tomé grosbeak *Crithagra concolor* and the abundant Príncipe seedeater *C. rufobrunnea* (Fig. 1). The latter species also occurs on two neighbouring islands, Príncipe and Boné de Jóquei (Fig. 1). Until recently, the grosbeak was placed in a monotypic finch genus and this incorrect taxonomy caused the species pair to be overlooked in a global analysis of congeneric bird taxa inhabiting small oceanic islands ^11^. Our previous work confirmed that the São Tomé grosbeak is indeed a gigantic seedeater with large bill and body, related to the canaries of the *Crithagra* clade, and a sister species to the more average-sized Príncipe seedeater^13^. To date, the process by which they speciated remains unknown.

**Figure 1.**
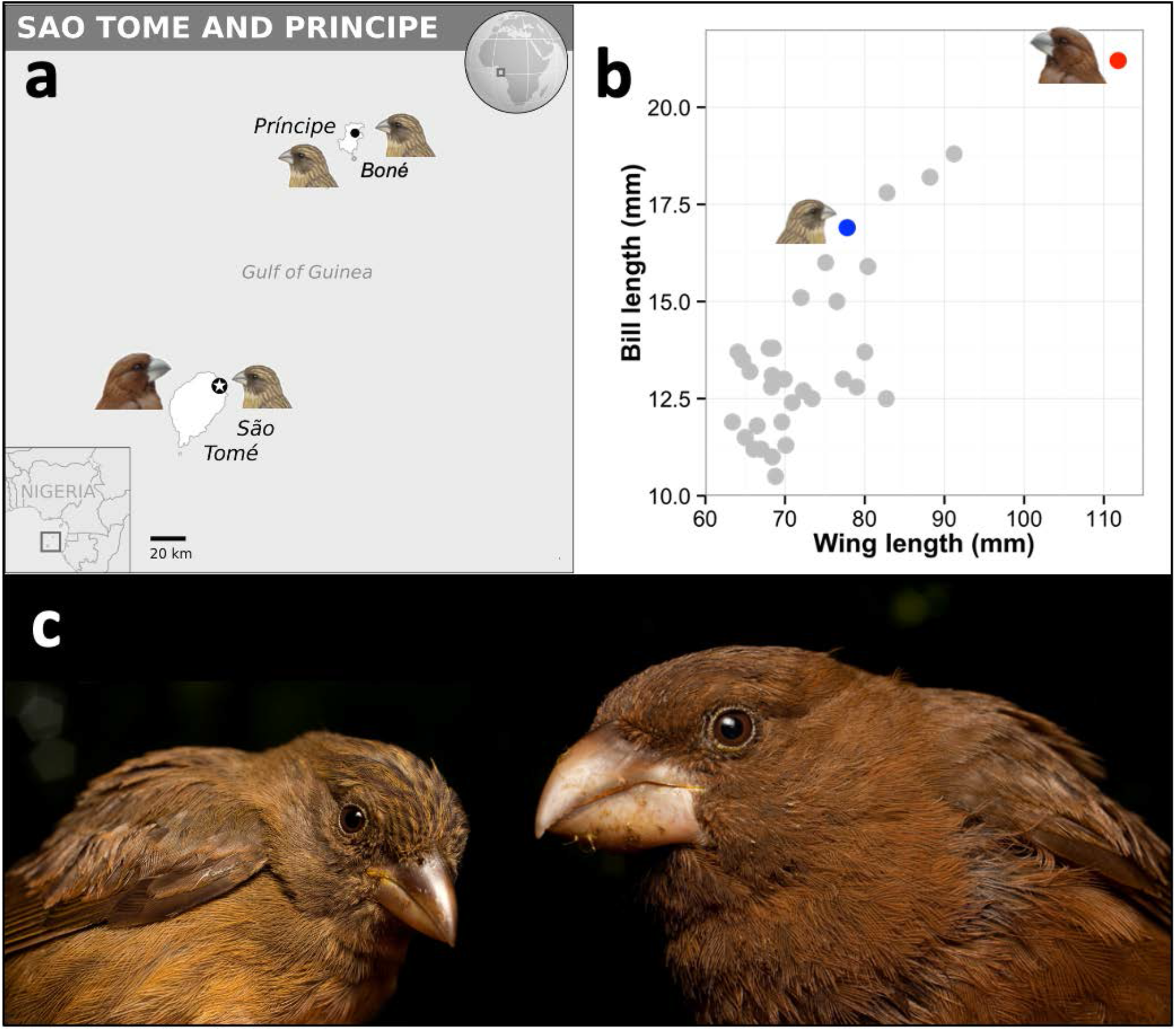
Geographic distribution and relative bill and body size of the São Tomé grosbeak and the Príncipe seedeater. **(a)** The São Tomé & Príncipe islands are situated in the Gulf of Guinea, Central Africa. The Príncipe seedeater *Crithagra rufobrunnea* occurs on Príncipe, the islet Boné de Jóquei, and São Tomé. The São Tomé grosbeak *C. concolor* is restricted to São Tomé, where it is sympatric with the local Príncipe seedeater population. **(b)** Wing length and bill length in males among species in the genus *Crithagra*. The Príncipe seedeater is highlighted in blue and the São Tomé grosbeak in red. **(c)** The Príncipe seedeater (left) and São Tomé grosbeak (right). Figure modified, with permission from the BOU/Wiley, from Melo et al. (2017). Photos: Alexandre Vaz. Map adapted from UN Office for the Coordination of Humanitarian Affairs (OCHA). Illustrations by Hilary Burn from the Handbook of the Birds of the World, with permission from Lynx Edicions.

## RESULTS

### Unresolved phylogenetic placement of the São Tomé grosbeak

Phylogenetic analyses of the São Tomé grosbeak and the three populations of the Príncipe seedeater, based on multiple genetic markers (sequences of 2 mitochondrial DNA [mtDNA] and 33 nuclear introns and exons; 34 microsatellites; see SI) and using both concatenated and species tree approaches, placed the São Tomé population of the Príncipe seedeater as a sister taxon to the São Tomé grosbeak, rather than to its two conspecific populations on Príncipe and Boné de Jóquei (Fig. S1–3). The same topology was obtained for the species tree inferred from 9,892 well-spaced single nucleotide polymorphisms (SNPs), recovered through Restriction site-Associated DNA (RAD) sequencing (Fig. 2a). These results are in line with sympatric divergence of the grosbeak and the seedeater population on São Tomé. However, gene flow between incipient species can have pervasive effects on phylogenetic inference and make it difficult to distinguish between divergence in sympatry and introgression upon secondary contact using phylogenetic approaches^14-16^. In line with this reasoning, a Bayesian species tree approach based on the mtDNA and nuclear markers implemented in StarBEAST gave equivocal results regarding the phylogenetic placement of the grosbeak: 75% of the posterior distribution of species tree topologies favoured a monophyletic seedeater clade (Fig. 2b, red tree set), whereas only 19% grouped the grosbeak with the seedeater population on São Tomé (Fig. 2b, blue tree set).

**Figure 2.**
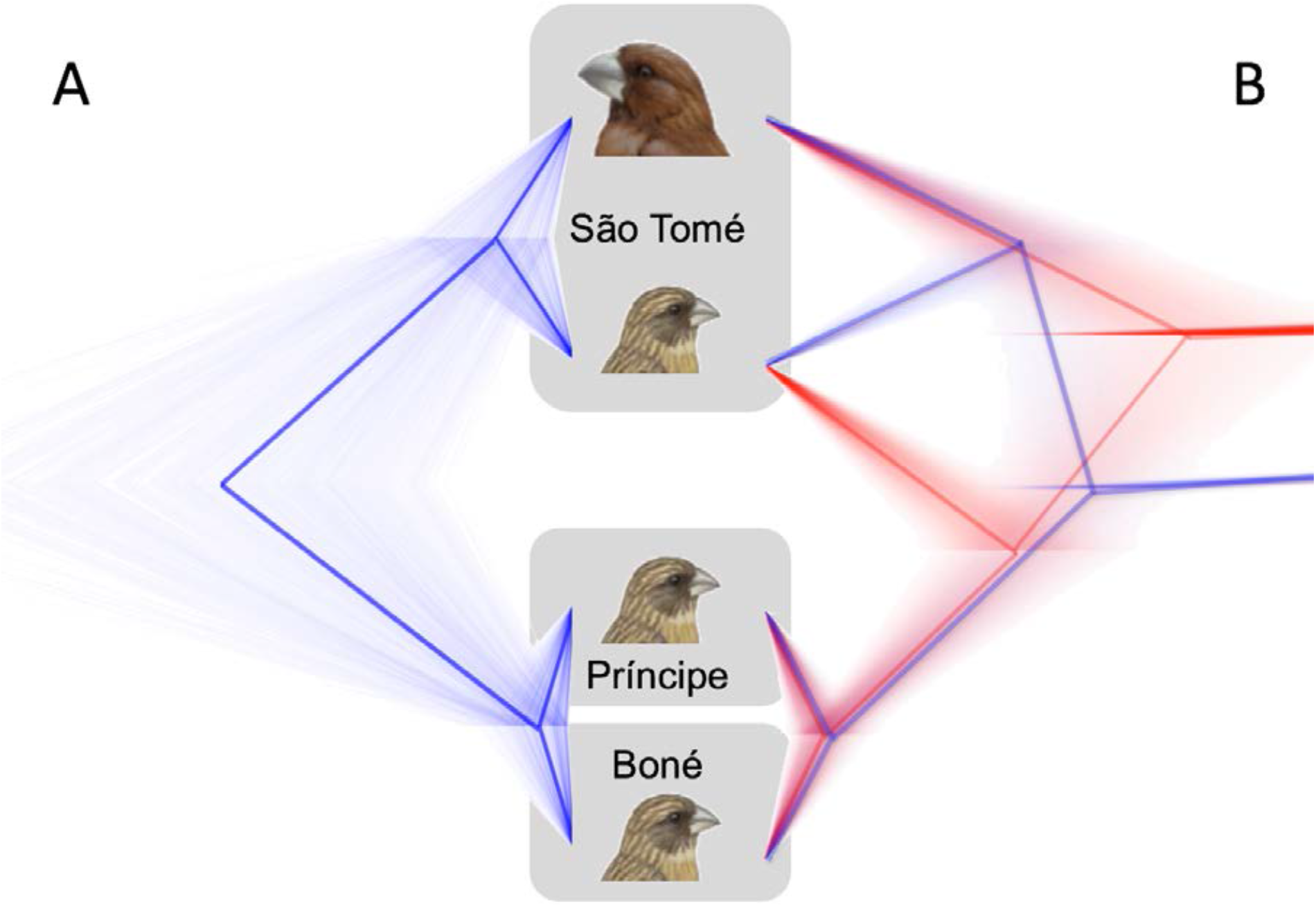
Different coalescence analyses result in conflicting relationships between the São Tomé grosbeak and the three populations of its sister species, the Príncipe seedeater. **(a)** Analyses of 9,892 SNPs evenly distributed over the genome in SNAPP resulted in a topology that groups the sympatric populations of São Tomé grosbeak *Crithagra concolor* and Príncipe seedeater *C. rufobrunnea* on São Tome, concordant with speciation in sympatry. **(b)** Analyses of 2 mitochondrial and 31 nuclear sequence markers in StarBEAST rendered two major topologies: 19% of the posterior distribution of species tree topologies (blue tree set) corresponded to the topology recovered by SNAPP, while 75% of the trees (red tree set) placed the grosbeak basal to all seedeater populations. A third topology (not shown) corresponding to 6% of the trees, groups the São Tomé grosbeak and the Príncipe and Boné de Jóquei populations of the Príncipe seedeater. The topology represented by blue trees was also recovered in other phylogenetic analyses using mitochondrial and nuclear sequences and in phenetic analyses using microsatellite genotypes (see SI).

### Genome-wide allele data reveal isolation and introgressive hybridization

To distinguish between divergence in sympatry and introgression upon secondary contact between previously allopatric populations, we analysed the full RAD sequencing dataset comprising 132,000 biallelic SNPs. These data showed two compelling results. First, an analysis of all SNPs where one allele was fixed in one population whereas the alternative allele was fixed in the other three populations, showed that the grosbeak harboured 1,193 private alleles, whereas each seedeater population had fewer than a hundred private alleles (São Tomé 82, Príncipe 74 and Boné 8; Fig. 3). Second, at all SNPs where one allele was fixed in two populations whereas the alternative allele was fixed in the other two, the grosbeak shared many more alleles (882) with the seedeater population on São Tomé, than it did with the seedeater populations on Príncipe and Boné de Jóquei (0 and 1, respectively; Fig. 3). These results strongly suggest that the grosbeak accumulated many unique mutations (private alleles) during periods of isolation, some of which introgressed into the seedeater population that colonised São Tomé. The alternative scenario with speciation in full sympatry would have required the grosbeak to have a 14-fold higher rate of molecular evolution in comparison to its sister species (1,193 vs. 82 private alleles). Adding a mainland outgroup (*C. burtoni*) to the analyses, ABBA-BABA tests confirmed introgression with a *d* statistic of 0.2 (where 0.0 signifies no introgression and 1.0 complete introgression) that was highly significant (Z score >18.5; Table S1). Based on this pattern of allele sharing and introgression between São Tomé populations (Fig. 3) and the current geographic distribution of the two species (Fig. 1), the most parsimonious demographic inference is that the grosbeak occurred in isolation on São Tomé until the seedeater colonized São Tomé from Príncipe.

**Figure 3.**
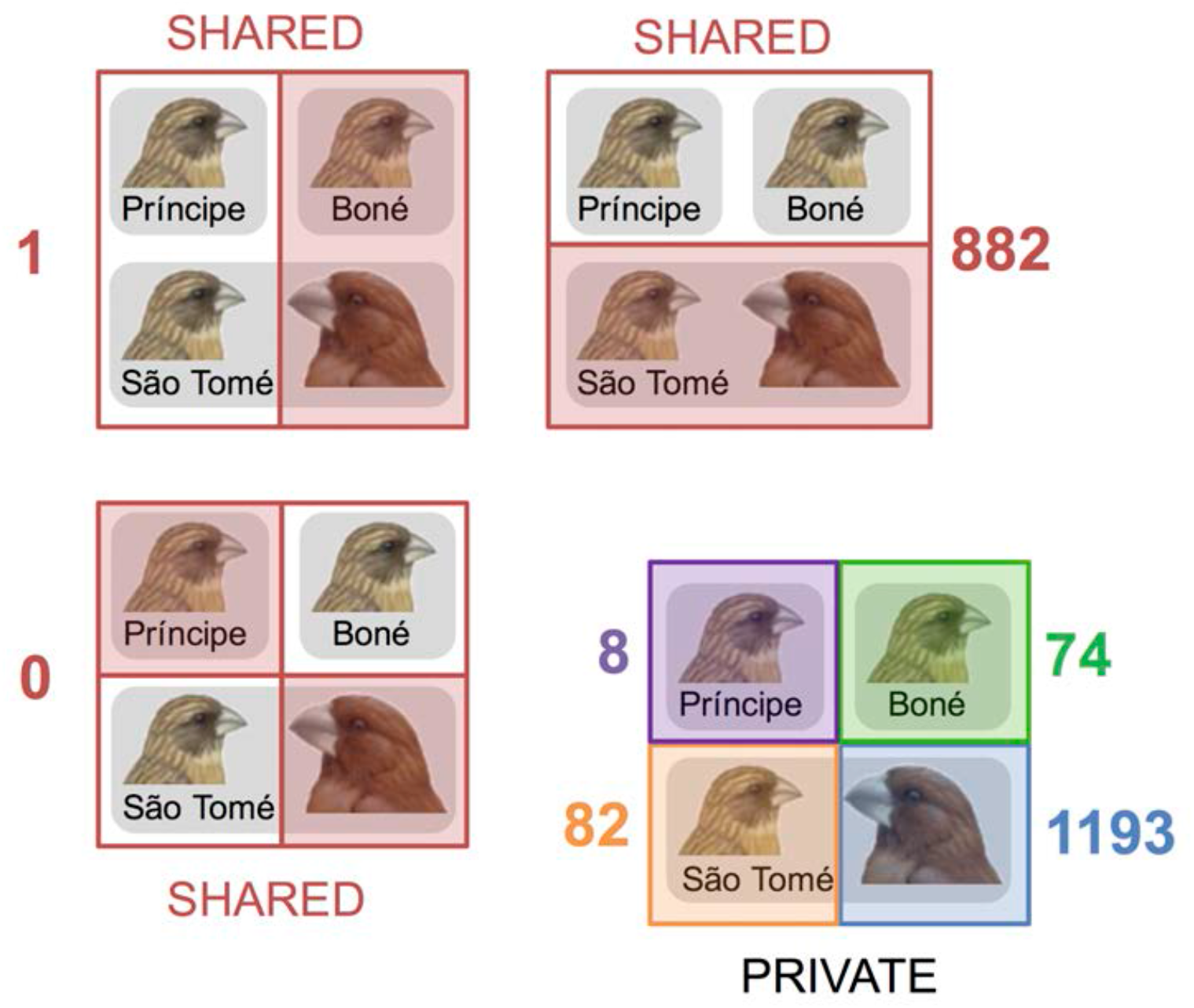
Distribution of private and shared alleles in the São Tomé grosbeak and the three Príncipe seedeater populations. Three panels (red font and rectangles/squares) show the number of biallelic single nucleotide polymorphisms (SNPs) where the same allele (e.g., the ‘red’ allele) was fixed in two populations whereas the alternative allele (e.g., the ‘white’ allele) was fixed in the other two. For the three pairwise combinations, the São Tomé grosbeak *Crithagra concolor* shares many more alleles with the sympatric population of the Príncipe seedeater *C. rufobrunnea* on São Tomé (882), than it does with the seedeater populations on Príncipe and Boné de Jóquei (0 and 1, respectively). The lower right panel shows the count of SNPs where one allele was fixed in a single population whereas the alternative allele was fixed in the other three populations (‘private alleles’). Notably, 1,193 fixed alleles are unique to the grosbeak, compared to only 82 unique fixed alleles in the largest population of the seedeater.

During secondary contact, introgression might either be asymmetric (unidirectional) or symmetric (bidirectional). Our SNP allele-sharing analysis (above) is inherently biased towards detecting introgression from the diverging grosbeak population into the seedeater population upon secondary contact, since introgression from the seedeater to the grosbeak creates a situation where the grosbeak will share alleles with all three seedeater populations, removing detectable variation. However, isolation-with-migration (IMa) models based on the nuclear sequences (33 loci) did also show evidence of gene flow from the seedeater population on São Tomé to the grosbeak (Table S2). This suggests that the grosbeak population did receive genetic material after secondary contact. The IMa models further estimated the split between the grosbeak and seedeater populations on São Tomé to have occurred ∼0.6 million years ago (Mya; Table S2). This is similar to the divergence times estimated with BEAST, which placed the splitting of the allopatric populations of Príncipe seedeater, and the split between the sympatric seedeater and grosbeak populations (i.e., the time of the last major introgression event), at ∼1.0 Mya and ∼0.7 Mya, respectively (Table S3).

### Non-introgressed genome regions support selection during divergence

Next we analysed the phylogenetic signal, and the nucleotide diversity and genetic differentiation, in local regions of the genome using the RAD sequence data and by using the zebra finch *Taeniopygia guttata* genome as a reference (taeGut3.2.4; Warren et al. 2005). Cross-species synteny mapping is feasible in passerines owing to a remarkably conserved karyotype among many bird species^17,18^. These analyses showed that introgressive hybridization between the grosbeak and the seedeater population on São Tomé has resulted in a mosaic pattern of divergence and introgression over the genomes (Fig. 4, S4, S5).

**Figure 4.**
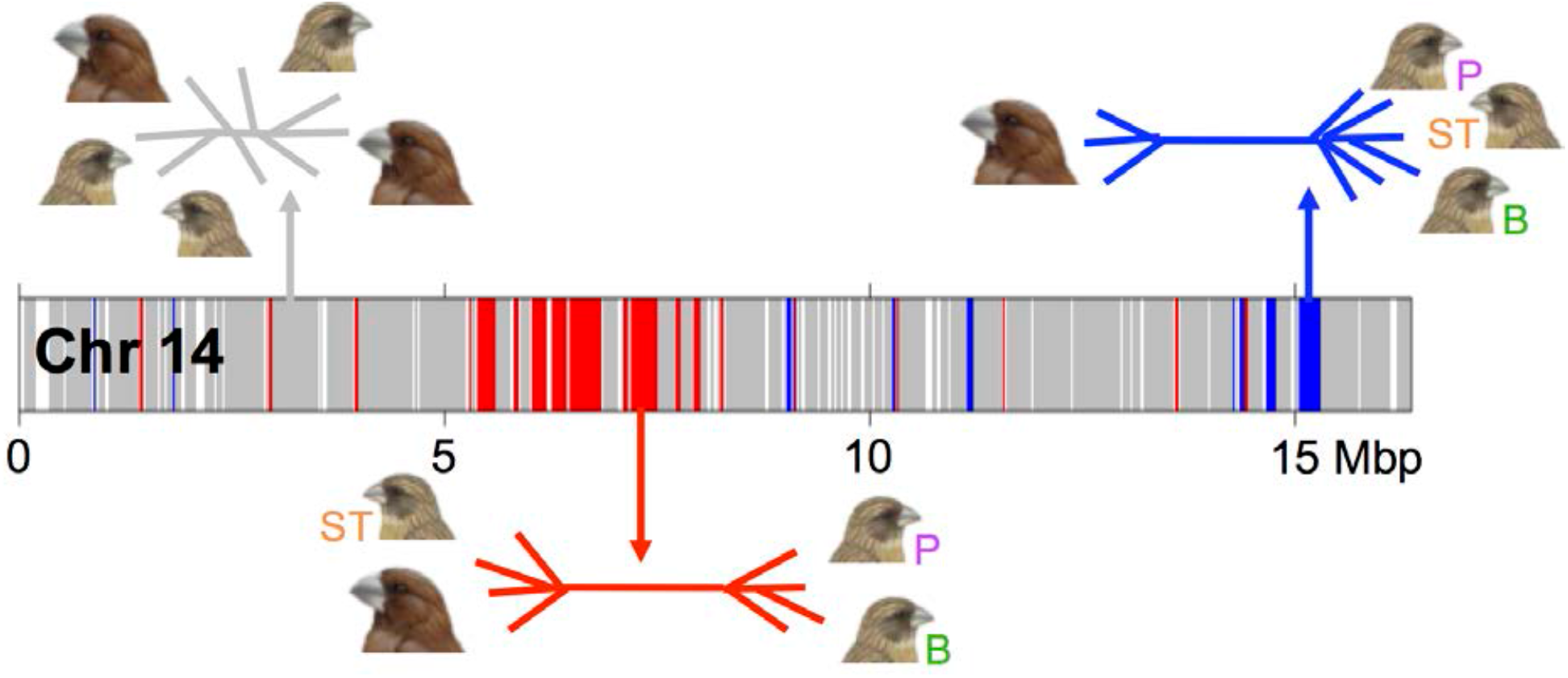
Genomic patterns of differentiation and introgression between the São Tomé grosbeak and the Príncipe seedeater. exemplified by data from chromosome 14. Three classes of phylogenetic signatures are distributed across the genome: Chromosomal segments that are phylogenetically inconclusive are grey (‘inconclusive’; 90.7% of all segments across the genome that were assigned a local phylogeny). Segments representing a preserved phylogenetic signal of the São Tomé grosbeak *Crithagra concolor* (not introgressed during secondary contact), where the three Príncipe seedeater *C. rufobrunnea* populations are sister taxa separated from the grosbeak, are blue (‘preserved’; 4.6%). Segments representing introgression from the grosbeak to the São Tomé population of Príncipe seedeater are red (‘introgressed’; 6.3%). This renders the sympatric populations of the grosbeak and the seedeater on São Tomé as sister taxa, separated from the other seedeater populations. Example topologies (‘cacti’) representative of each of the three genomic classes are drawn in corresponding colors, with the three seedeater populations abbreviated as ST (São Tomé; orange font; sympatric with the grosbeak), P (Príncipe; purple), and B (Boné de Jóquei; green). An extended version of this figure, including F_ST_ and nucleotide diversity (π) plots, is available as Figure S5.

Additional analyses of the mosaic genomes of the grosbeak–seedeater species pair on São Tomé provided support for selection against erosion of adaptive variation, i.e. the genomic segments that were not introgressed in either direction, and thus have been preserved with a unique genotype for the grosbeak (‘preserved’; see Fig. 4), were located significantly more closely to genes than was the case for introgressed segments or segments of inconclusive phylogenetic signal. We tested this by dividing the intergenic spaces, which vary considerably in size, into 2×50 equally sized bins ranging from just adjacent to a gene (relative distance 0.00) to the point halfway to the next gene (relative distance 0.50), and counted the number of intersecting genomic segments classified as being either ‘preserved’, ‘introgressed’ or ‘inconclusive’. If the genomic segment classes show no spatial association to genes, one would expect them at an even frequency in each bin across the relative intergenic distance. All three genomic segment classes were associated with genes, i.e. more segments were close to genes than far from them, but this relationship was significantly stronger for the segments that were preserved from introgression in the grosbeak (Fig. 5; ANCOVA: frequency of segments ∼ relative bin distance to gene×class of phylogenetic signal; F_5,144_ = 8.56, adjusted r^2^ = 0.20, p < 0.001; effect of class [‘preserved’ different from ‘inconclusive’ and ‘introgressed’] b = 0.001, p = 0.030; effect of relative distance to gene b = – 0.002, p = 0.068; effect of interaction of distance and class [‘preserved’] b = –0.004, p = 0.011). Note that the general association of the genomic segments to genes is likely owing to methodological issues, including non-random distribution of restriction enzyme recognition sites (and hence RAD sequencing markers) and lower interspecific mapping success in non-genic regions, leading to fewer genomic segments associating with bins far from genes.

**Figure 5.**
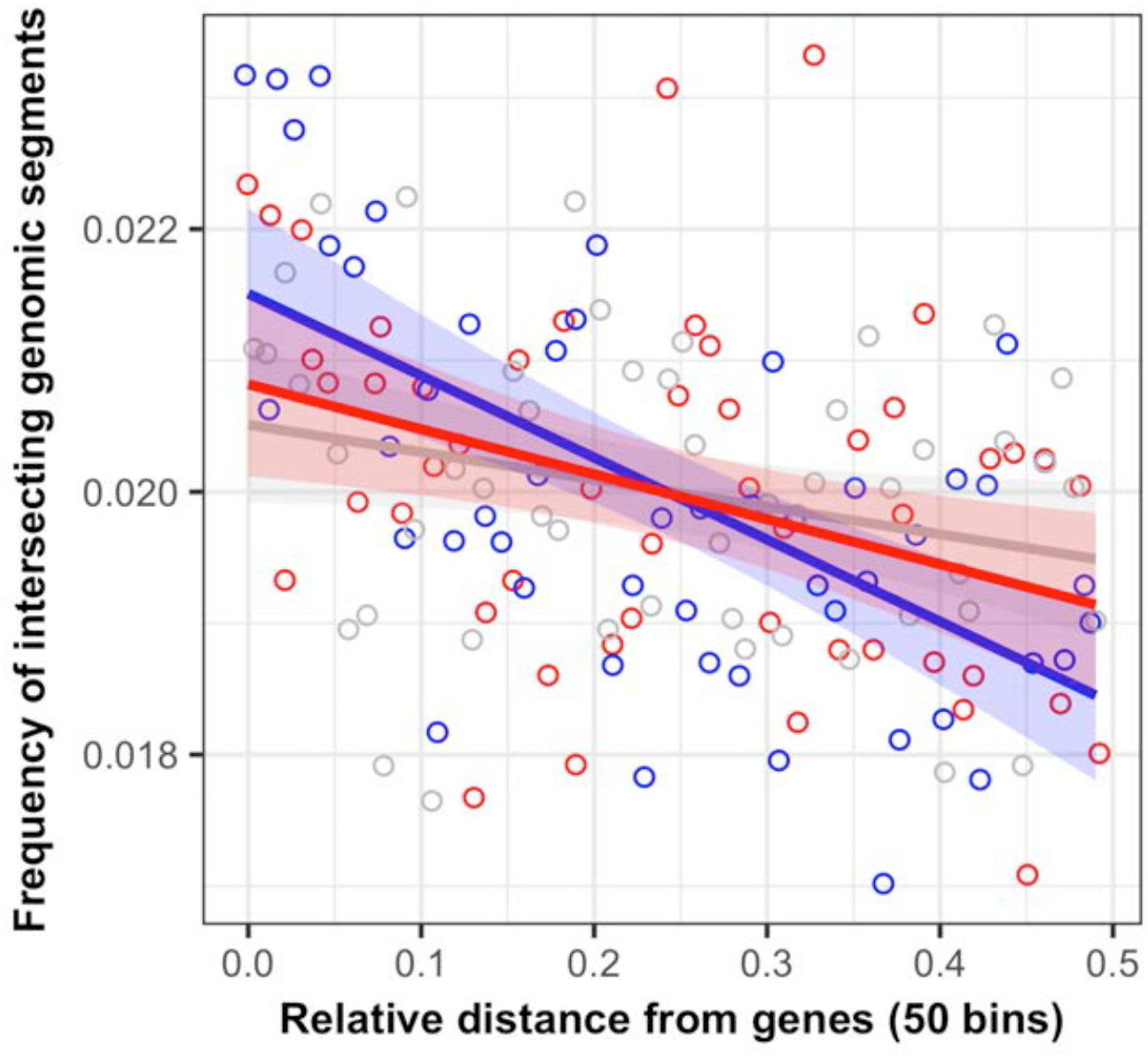
Proximity to genes varies between classes of genomic segments with different evolutionary histories. (colours correspond to Figure 4). The distances between genes vary and were normalized to a relative distance where 0 is immediately next to a gene and 0.5 is halfway to the neighbouring gene. In general, more genomic segments are found in bins near genes, and fewer in the bins further from genes (see main text). However, the strength of this association differs significantly between genomic segment classes (i.e., slopes differ significantly from each other, se ANCOVA statistics in main text), where non-introgressed (‘preserved’) segments (blue), containing alleles unique to the São Tomé grosbeak *Crithagra concolor* and different from all Príncipe seedeater *C. rufobrunnea* populations, are more strongly spatially associated to genes than the two other classes. Values from segments that are phylogenetically inconclusive are grey (‘inconclusive’) and segments representing introgression from the grosbeak to the seedeater population on São Tomé are red (‘introgressed’). Linear regression lines with translucent 95% confidence intervals are shown.

Furthermore, among 22 genes whose effect on bill morphology in birds has been established (Table S4), five (23%) overlapped with six ‘preserved’ genomic segments directly or within 5 kbp upstream/downstream of the gene, where regulatory elements may be found. The five recovered genes determine bill length, depth, and width: the bone morphogenetic protein 2 (BMP2^19^), the transforming growth factor beta receptor II (TGFBR2^20-22^), the high mobility group AT-hook 2 (HMGA2^23,24^), the collagen type XXVII alpha 1 chain (COL27A1^25^), and the fibroblast growth factor 8 (FGF8^21,26^). The ‘preserved’ segments make up only 48.6 Mbp (3.9%) of the reference genome assembly (taeGut3.2.4). Through bootstrapping, we shuffled genomic ranges corresponding exactly to the sizes of all ‘preserved’ genomic segments, across the genome 10,000 times and explored their intersection with the 22 bill genes (including upstream and downstream regions). Only in 1.2% of these bootstraps did six or more randomized genomic segments overlap with (any number of) bill genes, and only in 3.8% did (any number of) randomized genomic segments overlap with five or more bill genes. Since body size is highly polygenic with small effect sizes^25,27^ it cannot be analysed in the same way.

## DISCUSSION

Our genetic data suggest that speciation in the São Tomé grosbeak–Príncipe seedeater system involved a period of isolation on São Tomé, during which the grosbeak lineage acquired mutations through genetic drift and natural selection. These are detectable as private SNPs in the present-day grosbeak population and as shared alleles in the co-occurring grosbeak–seedeater populations on São Tomé. It is likely that the grosbeak started to evolve a large body and bill size during an initial period in isolation, a well-documented evolutionary trend in small birds following the colonization of oceanic islands^28-34^. Several factors may explain this trend, most of which linked to the low levels of interspecific competition (and of predation, often absent) and correspondingly high levels of intraspecific competition that characterize species-poor island communities^31,35^. Interestingly, although lack of interspecific competition may have been one component setting the grosbeak on the path of increasing body and bill size, the ecological factor that may have driven this species all the way towards ‘gigantism’—the grosbeak is the largest seedeater in the world (Fig. 1b; Melo, et al.^13^)—is likely to be the emergence of interspecific competition after the colonization of São Tomé by the lineage that evolved into the Príncipe seedeater. The co-occurrence between the two closely related lineages is expected to have led to intense competition further promoting the divergence of the grosbeak away from the typical seedeater size (character displacement; Losos^36^). This hypothesis was suggested seventy years ago^37^ to explain the origin of the three ‘giant’ birds from São Tomé: the grosbeak, the giant weaver *Ploceus grandis*, and the giant sunbird *Dreptes thomensis*. Our results show a clear and extensive genomic signature of introgressive hybridization between the grosbeak and the seedeater population on São Tomé. Introgression probably took place in both directions, and gene flow from the seedeater into the grosbeak lineage may have allowed the latter to restore genetic variation (cf. Table S5). However, importantly, our results show that selected parts of the genome including significant enrichment of genes related to bill size and shape, remained unique to the grosbeak. This supports the suggestion that resource-driven selection has been implicated in the evolution of the grosbeak’s extreme size.

A period of isolation was probably crucial for speciation because it would have created favourable conditions for the speciation process to be completed when the two closely related lineages met on São Tomé. Mutations accumulating in each population, through selection or genetic drift, may have created intrinsic incompatibilities, such as Bateson-Dobzhansky-Muller incompatibilities^38^. Incompatible loci strengthen the coupling of individual genes spread across the genome and can lead to the establishment of linkage disequilibrium between adaptive and assortative mating traits^14^. Nevertheless, the grosbeak–seedeater lineages that met on São Tomé hybridized and exchanged genetic material, suggesting that any incompatibilities that may have evolved in allopatry had not yet caused complete barriers to gene flow. The build-up of some phenotypic divergence in isolation could have facilitated the speciation process upon secondary contact (cf. Pfennig and Pfennig^39^) by decreasing the likelihood of competitive exclusion of one of the lineages, and by providing variation in traits that could be co-opted for assortative mating. Body and bill size are traits known to play a major role in bird feeding ecology and niche separation— and hence in species co-existence in sympatry^29,40,41^. The substantial size difference between the grosbeak and the seedeater would constitute an effective reproductive barrier important for completing the speciation process, as has been documented also for other systems^1,2,8^. For example, in Darwin’s finches, body size differences appear to be the most effective barrier to gene flow, independently of song differences and of phylogenetic relatedness^8^. Moreover, theory and data have shown that divergence with gene flow is facilitated when selection for reduction of gene flow acts on a large part of the genome, rather than being dependent on selection on a few loci^14,15,42^. Such a situation is highly likely in the grosbeak–seedeater system as the traits under selection (bill size and shape, and body size) are well known to be under polygenic control^21,43,44^. This is supported by the relatively high level of unique polymorphisms in the grosbeak (Fig. 3) and the genome-wide signal of selection (Fig. 5).

In conclusion, our genomic data suggest that the São Tomé grosbeak population has diverged partly in isolation, that the grosbeak–seedeater lineages hybridized after coming into contact causing extensive genomic introgression, and that resource-driven selection drove ecological divergence and turned one of the resulting species into an island giant. Thus, we provide a rare demonstration of isolation, hybridization and selection acting in concert during the speciation process.

## METHODS

### Sample collection and DNA extraction

We collected blood samples non-destructively from mistnetted wild birds (Table S6) and extracted DNA using DNeasy Tissue Extraction Kits (Qiagen) or standard phenol-chloroform protocol^45^.

### Mitochondrial sequence markers

To clarify the relationships between the three allopatric populations of *C. rufobrunnea* and *C. concolor*, we sequenced 10 Príncipe seedeaters from São Tomé, 10 from Príncipe and 9 from Boné for the ATP8 and ATP6 mitochondrial genes, together with 3 São Tomé grosbeaks and non-sister mainland African outgroups (*C. albogularis, C. burtoni, C. striolata*, and *C. sulphurata*), following Melo, et al.^13^. We used primers and PCR conditions from Melo^46^ and purified PCR products with Qiagen PCR Purification Kit, and used amplification primers to sequence both DNA strands with dye-labelled termination on an ABI 3730 sequencer. We aligned sequences in Geneious 4.7 (https://www.geneious.com) and verified that no gaps, insertions or deletions (indels) were present, and that all sequences translated into amino acids. We found no indications of having amplified nuclear copies of mitochondrial genes.

### Phylogenetic analyses of mitochondrial sequence markers

Molecular phylogenies were estimated using maximum-parsimony as implemented in PAUP* v. 4.0b10^47^, and model-based approaches (maximum likelihood, ML and Bayesian inferences, BI), as implemented in RAxML version 2.4^48^ and MrBayes 3.1^49,50^. The most adequate partition of the sequence data together with the best substitution model for each partition were evaluated using PartitionFinder^51^. Each Bayesian search consisted of two independent runs of four incrementally heated Metropolis-coupled MCMC chains run for ten million generations with trees sampled every 1000 generations, with the first 25% discarded as ‘burn-in’ period. The log-likelihood values and posterior probabilities (PP) were checked to ascertain that the chains had reached stationarity, and the effective sample size (ESS) of the log-likelihoods was estimated in Tracer v. 1.5^52^ to confirm further that the parameter space was properly explored. The concatenated analyses were performed freeing the different parameters (base frequencies, rate matrix, shape parameter, proportion of invariable sites) to vary between the partitions (genes and codon positions). PP were estimated from the 50% majority-rule consensus tree of the 22,500 sampled trees. In RAxML, 10 independent thorough maximum-likelihood searches were performed and support was calculated from 100 bootstrap pseudoreplicates.

### Nuclear sequence markers

We selected 35 primer pairs amplifying intronic or exonic nuclear sequences in passerines^53-59^ and further designed 49 new primer pairs spread over different chromosomes by first aligning Expressed Sequence Tags (ESTs) from zebra finch with the chicken *Gallus gallus* genome build 2.1 in DIALIGN^60^. We selected ESTs that aligned with chicken DNA but exhibited gaps of 500–1,000 base pairs (bp), which we assumed indicated zebra finch introns. We used PrimaClade^61^ for designing degenerated primers in the exon regions, either flanking an intron or covering a single large exon.

We evaluated primers on 15 individuals of the passerine genera *Corvus, Acrocephalus, Parus, Taeniopygia, Crithagra*, and *Nesospiza*, optimizing a PCR protocol for 10 µl Qiagen Multiplex PCR Kit reactions containing 5 µl Qiagen Multiplex PCR Master Mix, 0.2 µl each of 10 µM forward and reverse primer, 2 µl template DNA (5 ng/µl), and 2.6 µl RNasefree water. We used a touchdown cycling protocol: activation at 95°C for 15 min; three step cycling with denaturation at 94°C for 30 s, annealing at varying temperatures (T_A_) for 90 s ^62^, and extension at 72°C for 90 s; final extension at 72°C for 10 min. We ran the protocol 35–40 cycles, lowering T_A_ by 0.5–1.0°C per cycle for the first 5–20 cycles (Table S7), setting initial T_A_ ≤1°C below primer melting temperature (T_M_) and final T_A_ 2–3°C below T_M_ (Table S7). PCR products were precipitated (NH_4_Ac and ethanol) and sequenced using amplification primers on an ABI Prism 3100 capillary sequencer (Applied Biosystems).

After evaluation of sequences^62^, 33 markers containing SNPs were used on an extended sample set of *C. rufobrunnea* and *C. concolor* (32 individuals) and non-sister outgroups (*C. burtoni* and *C. flaviventris*).

### Nuclear sequence editing, alignment, and haplotype inference

We trimmed sequences and searched for double peaks in the electropherograms using the Find Heterozygotes plug-in in Geneious, assigning them IUPAC codes. Sequence regions of too low quality, causing ambiguities, were coded as missing data. We aligned the sequences using the Geneious Align and MUSCLE Align options.

To obtain phased data for demographic analyses, we inferred haplotypes with the PHASE 2.1 algorithm^63,64^ available in DnaSP v. 5.0^65^. Given the close relationship^13^, we inferred haplotypes for *C. rufobrunnea* sspp. and *C. concolor* as one dataset, while haplotypes for each outgroup species were inferred separately. We ran the algorithm three times, with 100 burn-in iterations and 100 iterations (thinning interval 1), starting with seeds randomized within the intervals 0–20,000, 40,000–60,000, and 80,000–100,000, respectively. We used a model that allowed for recombination and set the initial estimate of the recombination parameter ρ^66^ to 0.000335, corresponding to an estimation of average population sizes based on Melo^46^. Suggested haplotypes for ingroup individuals were used for further analyses if genotype phase probability was ≥0.6, which corresponds to an estimated probability of correct phase call of 0.82 for this type of data^64^. If genotype phase probability was <0.6, we manually inspected whether a single site caused the low phase probability. If so, and if that site contained a unique mutation, the phased genotype was retained (meaning that, in on average 50% of the cases, the nucleotides of that mutation may have been assigned to the wrong allele). In other cases, the unresolved sites with nucleotide phase probability <0.6 were coded as missing data, whereas the haplotypes as whole could still be retained. For outgroups, we accepted all suggested haplotypes, despite often low phase probabilities (due to low sample sizes), since they would only serve as outgroups and not be in focus taxonomically.

### Concatenation analyses

We tested each of the nuclear markers (ingroup haplotype datasets) for selective neutrality using Tajima’s D calculations^67^ in DnaSP v. 5.0^65^ and for intra-locus recombination using (1) the R-statistics^68,69^ in DnaSP v. 5.0^65^ and (2) a Difference of Sums of Squares^70^ approach in TOPALi v. 2^71^, with a window size of 100 nucleotides and a step size of 10. For each marker, the most suitable model of nucleotide evolution (Table S8) was determined according to the Akaike Information Criterion (AIC) with MrModeltest v. 2.3 (https://github.com/nylander/MrModeltest2) and PAUP* v. 4.0b10^47^. Fixed indels within the ingroup were coded in separate partitions as restriction data. We concatenated datasets of (non-phased genotypic) sequence data containing (1) all available samples including partial sequences, or (2) a sample-restricted dataset including only complete sequences. For the sample-restricted concatenated dataset, *C. r. fradei* (samples SRF6 and SRF9), *C. r. rufobrunnea* (SRR1 and SRR3) and one *C. concolor* sample (NC3) were included although 1–2 small (non-overlapping) sequence fragments were unknown. As non-sister outgroup, we used either *C. burtoni* or *C. flaviventris* (depending on amplification success).

We analysed the above datasets in MrBayes 3.1.2^49,50^. For the sample-restricted concatenated dataset we ran four Markov chains for 5×10^6^ generations in two parallel replicates, with chain heating parameter set to 0.15. Trees were sampled at intervals of 1,000 generations, and PP were calculated from 2,500 trees after excluding ≥50% as burn-in. For the full concatenated dataset, the corresponding settings were 200×10^6^ generations, sampling interval 5,000, and 5,000 trees used for calculation of consensus and PP. We used Tracer v. 1.5^52^ to manually inspect plots of the likelihood scores, in order to ensure that they had reached stationarity.

### Coalescence-based analyses of Sanger sequencing markers

*Species tree analyses:* We used the largest non-recombining blocks of nuclear markers, and full mitochondrial markers, as input for *Beast^72^ in the Beast v. 2.1.3^73^ module BEAUti and implemented substitution model parameter estimates (except base frequencies) by MrModeltest. Clock models were linked for all nuclear markers, and tree models linked for the mitochondrial markers, whereas other tree and substitution models were left unlinked. We set the species tree prior to follow the Yule process and the population size model to follow the Piecewise linear with constant root. We ran seven independent MCMC runs with a chain length of 1×10^9^ generations, sampling every 10×10^5^ generation. The output was inspected with Tracer, in order to ensure that likelihood scores were stationary and that ESS were adequate. The different runs were then combined using the Beast component LogCombiner, after excluding 10% of each run as burn-in, and the trees were drawn using Densitree^74^. PP were computed using the Beast component TreeAnnotator.

#### Demographic analyses of nuclear markers

We used isolation-with-migration (IM) models^75^ implemented in IMa2^76^ to evaluate historical demographic parameters based on phased sequences of the *C. rufobrunnea* sspp. at Príncipe and São Tomé, and *C. concolor* as a sister clade. We did not include the subspecies *C. rufobrunnea fradei* at Boné de Jóquei islet because we wanted to keep the model simple, and because Boné was connected to Príncipe during glacial periods and as recently as 10 kya^77^. The modelled topology (i.e., the one most strongly supported by *Beast) for these three populations, was ((Príncipe, São Tomé), *C. concolor*). IMa2 estimates several demographic parameters, including population sizes (*Q*) of both extant and ancestral populations, split times (*t*) for the branching events, and asymmetric migration rates between populations (*m*_*aàb*_, *m*_*bàa*_). Applying the HKY substitution model, we modelled *Q* of the 3 extant (i.e. Príncipe, São Tomé and *C. concolor*) and 2 ancestral populations [i.e. (Príncipe, São Tomé), and ((Príncipe, São Tomé), *C. concolor*)]. Furthermore, we modelled *t* for both branching events, i.e. Príncipe–São Tomé (*t0*), and (Príncipe, São Tomé)–*C. concolor* (*t1*). Finally, we modelled *m* between the following population pairs: Príncipe–São Tomé, Príncipe–*C. concolor*, and São Tomé–*C. concolor*.

Upper bounds of prior distributions of parameter values were evaluated in several trial runs, and when PP peaks of all parameters fell well within prior boundaries, we ran two independent IMa2 runs with identical prior settings, but with different random seeds. The runs began with 10×10^6^ burn-in steps and then continued for >124×10^6^ steps, sampling every 1,000 genealogies. We achieved adequate convergence and mixing of the Markov chains as indicated by visual inspection of trend line plots, sufficient ESS and very similar parameter estimates and PP distributions in the two runs. We therefore averaged the values corresponding to the peak of the PP distribution as our parameter estimates. We calculated the effective population size (*Ne*) as *Q/*4*µk*, split times in years as *tg/µk*, and population migration rates per generation (2Nm) as *Qm/2*, where *µ* is the mutation rate per generation, *k* the mean length of the sequences (563.4 bp), and *g* the generation time (3 years^78,79^). The mutation rate was set to 4.6×10^−9^ substitutions per site per generation (ss^−1^g^−1 80^; cf. Bidegaray-Batista and Arnedo^81^). Note that the direction of gene flow for our *2Nm*-estimates are forward in time, i.e., *2N*_*C. concolor*_*m*_*São Tomé à C. concolor*_ indicates the number of migrants per generation that the *C. concolor* population receives from the São Tomé population.

### RAD sequencing

We prepared paired-end restriction site-associated DNA (RAD) libraries following Baird, et al.^82^ and Etter, et al.^83^, but applied no RNase A treatment, sheared DNA fragments with a Bioruptor Standard UCD-200 (Diagenode), and selected 150–750 bp fragments (excluding adapters). 50-µl PCRs were set up with 16 µl RAD library template running 16 cycles. We used restriction enzyme SbfI and multiplexed 16 samples per library, sequenced on one Illumina HiSeq 2000 lane each at BGI, Hong Kong. The paired raw reads were filtered with *process_radtags* (our barcodes differed by ≥ 3 nucleotides; --*barcode_dist* 3) and *clone_filter* in Stacks v. 1.21^84^. We mapped remaining reads to the zebra finch genome (taeGut3.2.4; http://genome.ucsc.edu/) with Bowtie v. 2.2.2^85^. Given the interspecific mapping we set maximum and minimum mismatch penalties (*--mp*) to 2 and 1, respectively. Further, we only accepted paired reads mapping in the expected relative direction within 150–1,000 bp from each other (*--no-discordant, --no-mixed*). The following bioinformatic handling followed two different tracks.

#### Stacks-based track

We used read 1 as input to Stacks, using *ref_map* for phylogenetic and coalescent analyses of the ingroup samples. We used default parameters except for increasing minimum within-sample sequencing coverage (m = 4). We then evaluated the ln likelihood scores of each catalogue locus with *rxstacks*, and set a cut-off at −238, which excluded 20% of the catalog loci that may have low coverage or high sequencing error rate. Confounded loci were also removed and excess haplotypes pruned in individuals.

We filtered the output to only contain loci which (1) had representation by ≥50% of samples for each population; (2) had been assigned 1–2 alleles in all samples; (3) contained 1–8 SNPs; (4) comprised ≤12 alleles, resulting in 42,424 loci. Stacks output was then converted to individual biallelic sequences, from which we defined a dataset where we accepted no missing data in any individual and required loci to be spaced by >5,000 bp, retaining 26,202 loci. These were concatenated based on sample (but with random allele pairing) to long super-matrices. Processing the alignment in VCFtools v. 0.1.12a^86^, we extracted a maximum of one SNP per RAD locus, with an overall minimum allele frequency of ≥0.10. Removing rare alleles is a necessity for efficient coalescence analyses with SNAPP^87^, since uninformative polymorphism can both slow and skew the analyses). The final dataset contained 13,909 SNPs. To further improve performance, we selected only 4 (biallelic) individuals from each population.

Using the Stacks *populations* we extracted all SNPs for which alleles were fixed within a population but differed between populations. As our sample size of *C. concolor* was lower than those of *C. rufobrunnea* sspp., we made one selection from the full dataset and one using four individuals from each population. We then tallied fixed private alleles and shared alleles between different populations.

#### Standard variant calling track

In a second round, we used the sample-specific BAM files from the mapping to zebra finch, sorted them with SAMtools v. 1.3.1^88^, added readgroup information (AddOrReplaceReadGroups) and merged them (MergeSamFiles) with PicardTools v. 2.8.1 (http://broadinstitute.github.io/picard), and then made variant calling using GATK v. 3.3.0^89^ and Samtools.

With GATK, we realigned indels with LeftAlignIndels, and made variant calls with UnifiedGenotyper^90^, allowing for maximum three alleles (one for the zebra finch reference, ≤2 alleles within our samples). We also called variants with SAMtools mpileup (flags –AEgf) and BCFtools v. 1.2^91^ (flags – vmO) and filtered the raw output with VCFtools v. 0.1.14^86^ minor allele count (--mac) at 1, to remove sites that were invariant within our samples but differed from the zebra finch reference.

GATK yielded 800,603 and SAMtools 1,062,989 SNPs. Using python, we extracted the intersecting 629,453 SNPs that were recovered using both variant callers. With VCFtools we extracted the ingroup (*C. concolor, C. rufobrunnea*) only, with 363,765 remaining SNPs, which were further quality filtered for SNPs genotyped in ≥75% of the samples at a sequencing depth of ≥8×, resulting in a final dataset of 131,661 SNPs.

### Coalescence-based species tree analyses of SNP data

The filtered SNP dataset from Stacks was imported into Beauti, where priors for forward (*u*) and reverse (*v*) mutation rates were set to be co-estimated during the runs. Remaining parameters were left at default values, i.e., species divergence rate λ = 0.00765, and θ defined by a γ prior with shape parameter α=11.750 and scale parameter β=109.73. The rate prior was set to follow an inverse gamma distribution. We performed two independent runs for ≥8.4M generations, sampling every 1K generation using the SNAPP v. 1.1.16^87^ plugin in Beast. Output was inspected with Tracer, and the converging runs were combined after excluding 10% as burn-in.

### Allele sharing and detection of introgression

Out of the 131,661 biallelic SNPs within the grosbeak– seedeater system, we selected those that were fixed for a single allele within each of the four population but differed between populations. We then tallied the cases in which one allele (e.g., *p* or the ‘red’ allele; cf. Fig. 3) was fixed in two populations whereas the alternative allele (e.g., *q* or the ‘white’ allele; cf. Fig. 3) was fixed in the other two populations. Further, we tallied cases in which one allele was fixed in a single population (a private allele) whereas all three other populations were fixed for the alternative allele.

Using the corresponding dataset, but with *C. burtoni* as outgroup (130,573 SNPs), we also performed D-statistic analyses on the same dataset using ABBA-BABA tests in Admixtools^92^ implemented in admixr^93^. We tested both the Príncipe and the Boné de Jóquei populations as sister to the São Tomé population of *C. rufobrunnea*. (Table S1).

### Detection of introgressed and preserved genomic regions, and their location relative to genes

We fed the 131,661 ingroup SNPs from GATK and Samtools into Saguaro (Zamani *et al*. 2013), which combines a Hidden Markov Model and a Self-Organizing Map to estimate local unrooted phylogenies (‘cacti’) along the chromosomes, without *a priori* information. After exploration of the effect of number of iterations, we ran 40 iterations, and filtered out cacti that eventually were not assigned to any genome part. We then collapsed the individual cacti into clustered classes, based on similarity, and recovered two distinct patterns (‘introgressed’ and ‘preserved’) and three classes that were inconclusive in terms of phylogenetic signal (Figures S4–5).

Using annotations of the zebra finch reference genome, we analysed spatial correlation between gene locations and genomic segments classified with different phylogenetic signals (‘introgressed’, ‘preserved’, or ‘inconclusive’) with the BEDTools v. 2.25.0^94^ command *reldist*. Since physical distances between neighboring genes vary, distances were normalized, and halfway intergenic spaces were divided into 50 equal bins in relative distance from genes. The intersections of those bins and the three genomic segments classes were counted and to test the potential association, we ran an ANCOVA with the *frequency of genomic segment class* as response variable, *relative distance to genes* as a covariate, and *class of phylogenetic signal* as a factor, including interactions.

We further analysed the intersection of ‘introgressed’ and ‘preserved’ genomic regions and candidate genes for bill morphology, based on literature review (Table S4). For genes HMGA2 and COL4A5 there have been major changes in annotation, including the range of the genes, in a later assembly (bTaeGut1_v1.p) and we therefore updated the start and end coordinates for those genes. We performed all analyses with BEDTools and included 5,000 bp upstream/downstream of the genes to cover any essential closely situated transcription factors. To ascertain whether the intersection with candidate genes was higher than expected at random, we assigned non-overlapping random parts of the genome exactly the sizes of the ‘introgressed’ and ‘preserved’ regions, respectively, and made 10,000 bootstrap replicates intersecting the actual genes. Results for ‘preserved’ regions are accounted for in the main text; ‘introgressed’ regions overlapped with two candidate bill genes, which does not represent any enrichment (57.3% of bootstrap replicates overlapped with ≥2 bill genes).

## Acknowledgements

We are thankful to everyone that made our work in the Gulf of Guinea possible. In São Tomé and Príncipe: A. Carvalho, S. Pontes, and V. Bonfim (Department of the Environment); C. Bom Jesus (Bikegila), P. Leitão, L. Mário, T. Pontynen, L. Primo, O. Veiga, P. Ryan, A. Thomasson, and A. Vaz (field assistance); the late A. Gascoigne for overall support. R. Covas was of invaluable assistance throughout. Fieldwork was supported by grants from the Genetics Society (UK) and the National Geographic/Waitt Foundation (W172-11). Parts of the analyses were carried out at the University of Oslo Bioportal (http://www.bioportal.uio.no/). This work was supported by the Swedish Research Council (621-2007-5381, 621-2009-4945, 621-2014-5222, 621-2016-00689), the Oscar and Lili Lamm Foundation, and the Crafoord Foundation, EU-FP7 (“Avian Genomics”) to B.H.; the Portuguese Science and Technology Foundation (PTDC/BIA-BEC/102253/2008, SFRH/BPD/46407/2008), the European Regional Development Fund (FCOMP-01-0124-FEDER-008934), and the North Portugal Regional Operational Programme 2007/2013 (“Genomics and Evolutionary Biology”) to M.M.; and the Royal Physiographic Society to M.S. Non-destructive sampling was carried out with the appropriate permissions from the Department of the Environment of São Tomé and Príncipe and the Ministry of Scientific Research and Innovation of Cameroon. Trevor Price, Peter Grant, Charlie Cornwallis, Rayna Bell, Mark Ravinet, and Miguel Carneiro kindly provided comments that have greatly improved this study.

## Author contributions

M.M. and P.J. conceived the study; M.S., M.M., and B.H. designed the study; M.M. performed most field work; M.S. and M.M. performed most lab work; M.S. and M.M. analysed most data, assisted by B.H.; M.M., M.S., and B.H. wrote the manuscript with input from P.J.

## Additional information

Additional material, and accession numbers for genetic material (Table S9), is available in the Supplementary Information. RAD sequencing data is available at the NCBI Sequence Read Archive under BioProject PRJNA792479: https://www.ncbi.nlm.nih.gov/Traces/study/?acc=PRJNA792479&o=acc_s%3Aa. Datasets are available at Zenodo: https://doi.org/10.5281/zenodo.5816808.

