## Supplementary Information for "Genomic signatures of isolation, hybridization, and selection during speciation of island finches"

#### SUPPLEMENTARY FIGURES

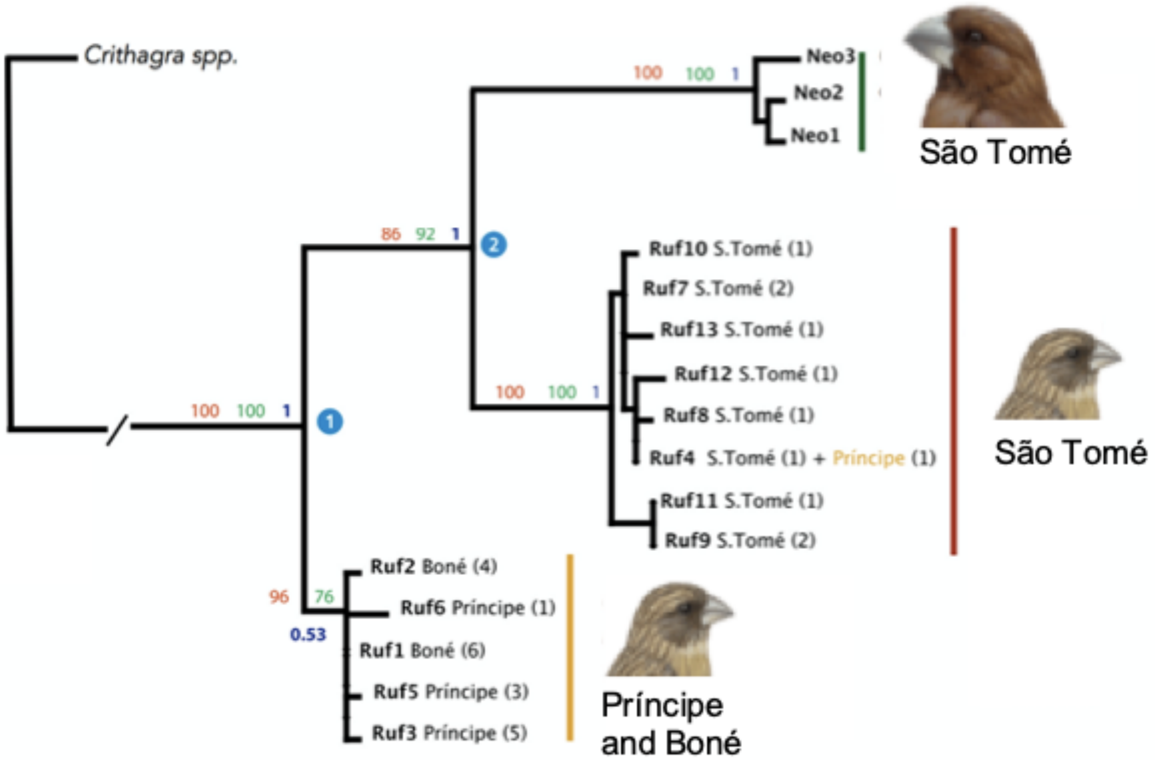

Figure S1. Phylogenetic tree of the São Tomé grosbeak *Crithagra concolor* (top; green clade) and the Príncipe seedeater *C. rufobrunnea* (bottom; red and orange clades) based on the ATP6 and ATP8 mitochondrial markers, with tRNA-Lys and fragments of COII. Mainland African outgroups were omitted for clarity. Support by maximum parsimony (red; 0–100), maximum likelihood (green; 0–100) and Bayesian inference (blue; 0–1) are stated at nodes. The two numbered nodes (white number in blue circle) indicate estimations of divergence times: (1) 0.97 million years ago (Mya), 95% confidence interval (CI) 0.59–1.42 Mya; (2) 0.73 Mya, 95% CI 0.41–1.07 Mya.

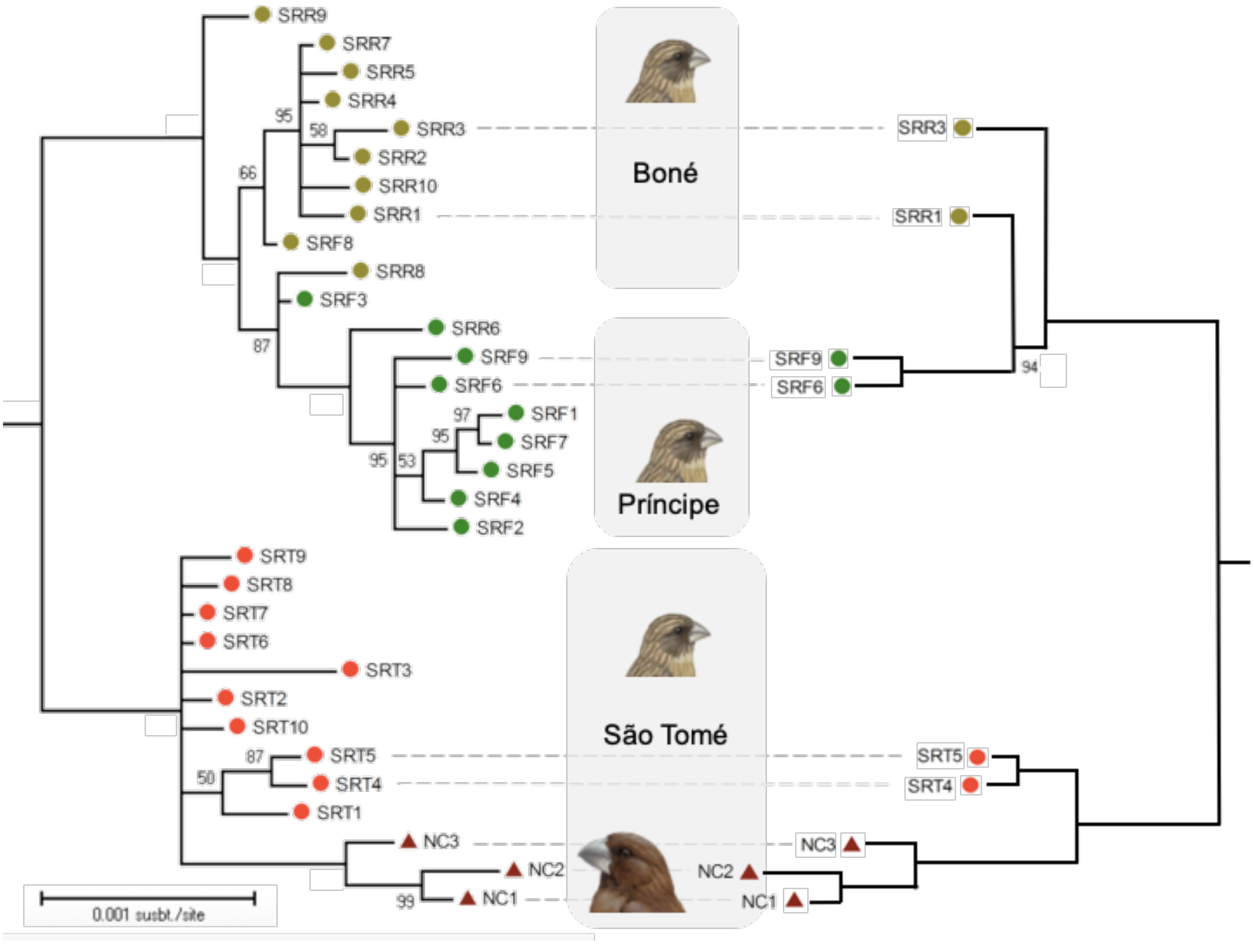

Figure S2. Phylogenetic tree of the Príncipe seedeater *C. rufobrunnea* (top; circles) and the São Tomé grosbeak *C. concolor* (bottom; dark red triangles) based on Bayesian inference of concatenated data from 33 nuclear sequence markers (>20 Kbp), computed with MrBayes. Seedeater populations are from Boné (light green circles), Príncipe (dark green circles), and São Tomé (light red circles). Mainland African outgroups *C. burtoni* and *C. flaviventris* were omitted from the figure for clarity. On the left side is a tree including all 33 markers and all 32 sampled ingroup individuals, that includes some missing data in the sequence matrix, as each marker contained 29–32 ingroup individuals with full or partial sequence data (mean 31.65 individuals per marker). Each of the 32 ingroup individuals is represented in 31–33 (mean 32.63) marker alignments with full or partial sequence data. On the right side is a tree with a subset of the sampled individuals, for which complete or very near complete sequence data was available for all 33 markers (allowing for 1–2 short partial gaps in five individuals, that did not colocalize between individuals). Dashed lines connect the same samples in the two datasets. Posterior probability (PP) is indicated as percentages, unless unlabeled, which indicates PP = 1.0 (100%).

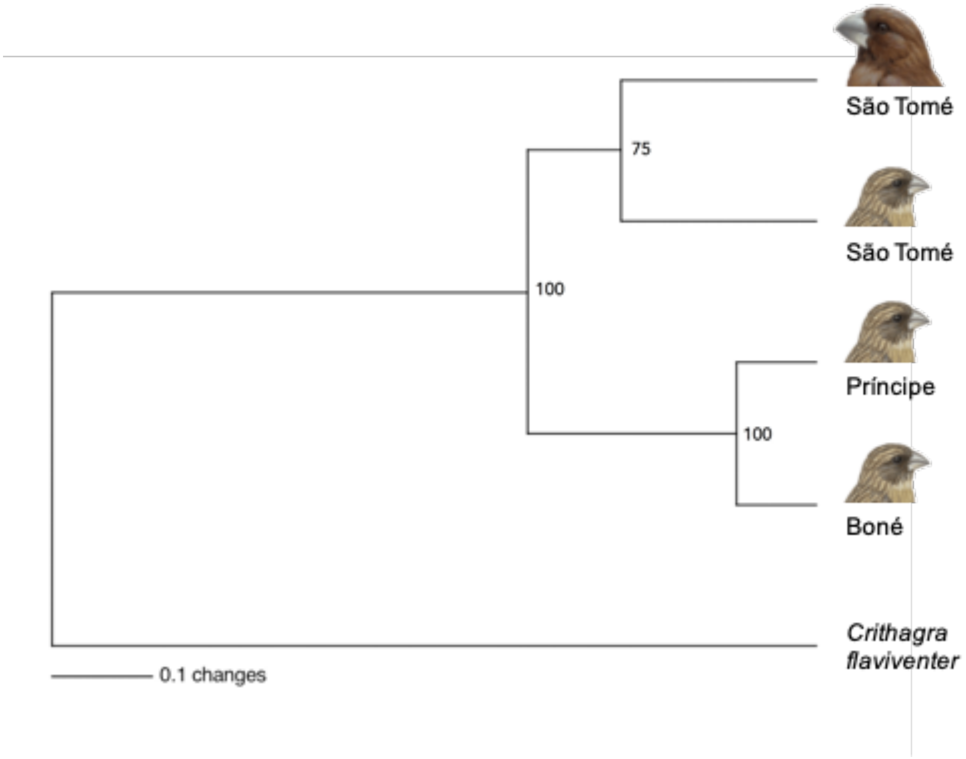

Figure S3. UPGMA population tree of Nei's distance, based on 34 microsatellites. The populations are São Tomé grosbeak *Crithagra concolor* on São Tomé (top; N = 3); the Príncipe seedeater *C. rufobrunnea* on São Tomé (N = 117), Príncipe (N = 37), and Boné de Jóquei (Boné; N = 67); with the mainland African yellow canary *C. flaviventris* as outgroup. Support values from 1000 bootstrap replicates are indicated as percentages at nodes. Cavalli-Sforza & Edwards's chord distance returned the same relationships (data not shown).

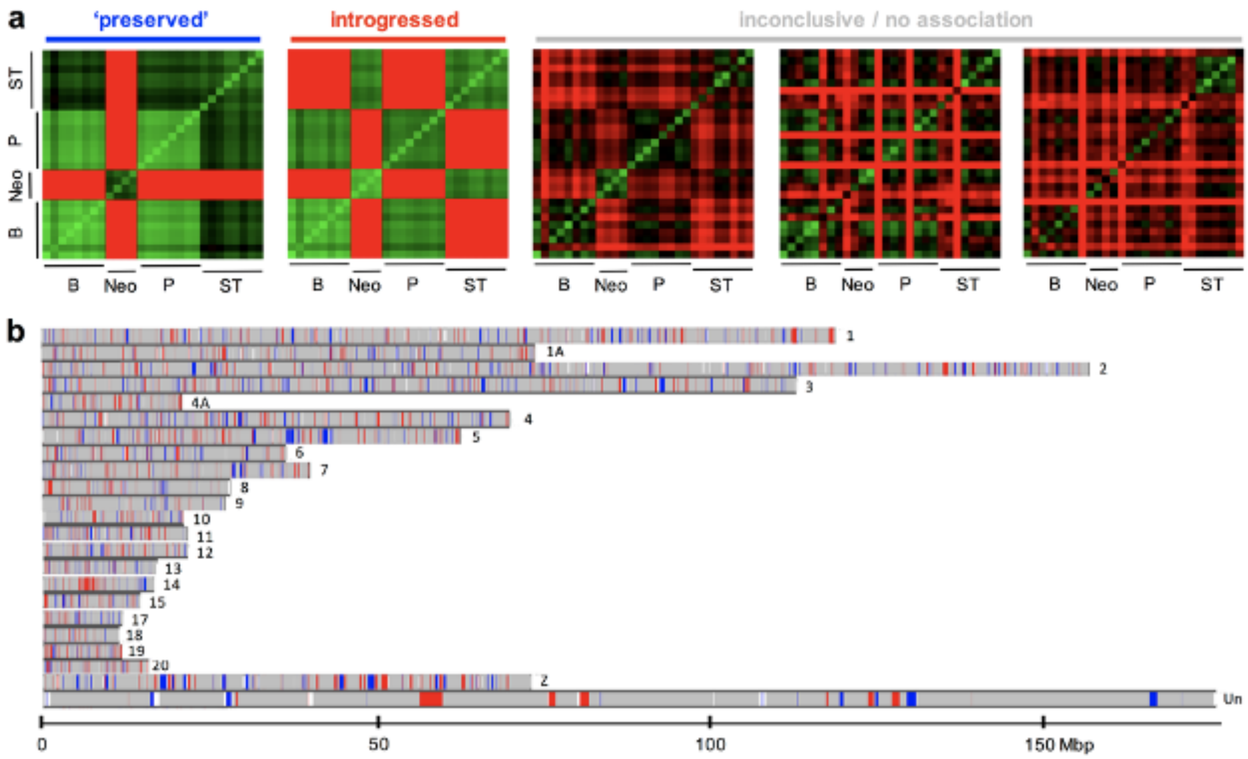

**Figure S4. (a)** Heatmaps of local phylogenies over the genome computed with Saguaro, collapsed into the five most representative topologies, representing three classes of phylogenetic signals ('preserved' from introgression with the São Tomé grosbeak *Crithagra concolor* differing from Príncipe Seedeater *C. rufobrunnea* populations – blue; introgressed from the grosbeak to the São Tomé seedeater population – red; inconclusive/no association – grey). Light green represents the highest similarity, which is lowered as the color darkens, and then turns red. The brightest red is the most dissimilar. Each cell represents one individual, and they are ordered along populations: Príncipe seedeater on Boné de Joquei (B), São Tomé grosbeak on São Tomé ("Neo"), Príncipe seedeater on Príncipe (P), and on São Tomé (ST). **(b)** The distribution of the three classes of phylogenetic signatures across the largest chromosomes (>10 Mbp) of the genome: phylogenetically inconclusive segments are grey (89.1% of all segments assigned a local phylogeny), segments with a 'preserved' grosbeak phylogenetic signal are blue (4.6%), and segments with a signal of introgression are red (6.3%).

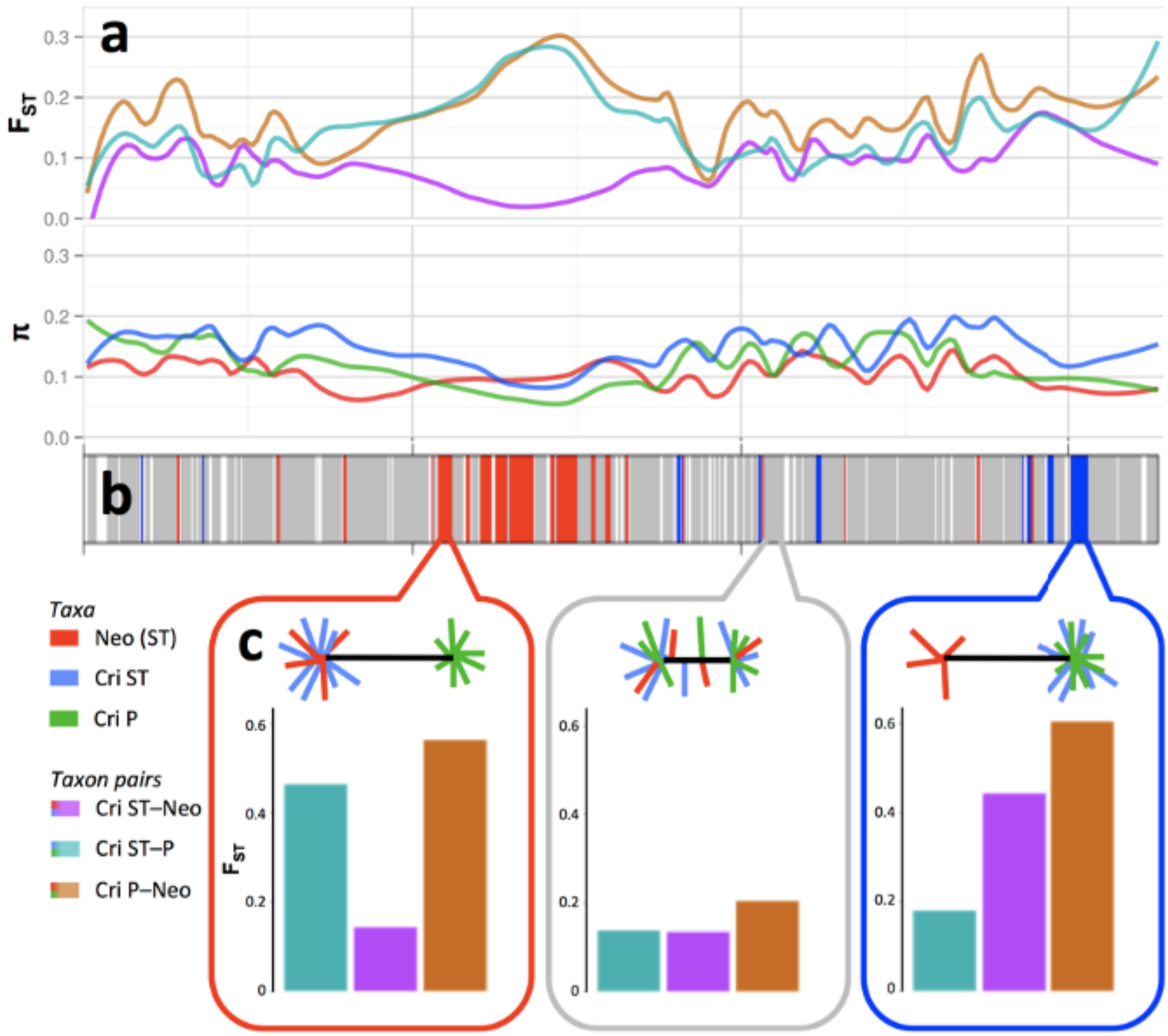

Figure S5. (a) Genomic patterns of differentiation and introgression between the São Tomé grosbeak *Crithagra concolor* and the Príncipe seedeater *C. rufobrunnea*, exemplified by chromosome 14. (a) FST and nucleotide diversity (π) in São Tomé grosbeak ("Neo"; red) on São Tomé (ST), the Príncipe seedeater (Cri) on São Tomé (blue), and on Príncipe (P; green). Colours for pairwise FST are mixes of the taxon-specific colours (see key). (b) Three classes of phylogenetic signatures across the genome: Chromosomal segments that are phylogenetically inconclusive are grey ('inconclusive'; 89.1% of all segments assigned a local phylogeny). Segments representing a preserved phylogenetic signal of the grosbeak (not introgressed during secondary contact), where the two seedeater populations are sister taxa, separated from the grosbeak, are blue ('preserved'; 4.6%). Segments representing introgression from the São Tomé Grosbeak to the São Tomé population of Príncipe seedeater are red. This renders the sympatric populations of the grosbeak and the seedeater on São Tomé as sister taxa, separated from the Príncipe population of Príncipe seedeater ('introgressed'; 6.3%). For the representation of other chromosomes, see Fig. S5. (c) Top panel: Examples of topologies ('cacti') representative of each of the three genomic classes described in b, colours as in a. Bottom panel: Genome-wide average FST for the three taxon pairs, per genomic class described in b (each box representing all segments in that class, over all chromosomes). Data for the seedeater population on Boné de Jóquei, which is genetically very similar to the Príncipe population, has been omitted for clarity.

### SUPPLEMENTARY TABLES

Table S1. D statistics from ABBA-BABA tests, evaluating introgression from the São Tomé grosbeak *Crithagra concolor* into the São Tomé (ST) population of Príncipe seedeater *C. rufobrunnea* using 130,573 SNPs. The mainland seedeater species *C. burtoni* is used as outgroup and Príncipe seedeater populations on (a) Príncipe (P) and (b) Boné de Jóquei (BdJ) as the sister to the São Tomé population of Príncipe seedeater.

| Topology | D statistic | Standard Error | Z score | N ABBA patterns | N BABA patterns |
|---|---|---|---|---|---|
| (((*C. rufobrunnea* P, *C. rufobrunnea* ST), *C. concolor*), *C. burtoni*) | –0.197 | 0.0104 | –18.9 | 1,713 | 2,555 |
| (((*C. rufobrunnea* BdJ, *C. rufobrunnea* ST), *C. concolor*), *C. burtoni*) | –0.209 | 0.0112 | –18.7 | 1,726 | 2,640 |

Table S2. Parameter estimates from IMa2 analyses of 28 non-recombining nuclear loci in three populations: the São Tomé grosbeak *Crithagra concolor* on São Tomé (N), and Príncipe seedeater *C. rufobrunnea* populations on Príncipe (P) and São Tomé (ST) (Boné de Jóquei population excluded). P+ST represent the ancestral population that split into the present populations of Príncipe seedeater on São Tomé and Príncipe; P+ST+N represent the mainland source population from which the islands were colonized. **(a)** Divergence time and population size estimates, in original units (t and q), and converted to years and individuals, respectively. **(b)** Migration rates, in original unit m and converted to number of migrants per generation. All conversions are based on mean sequence length (563 basepairs), an estimation of 4.6×10–9 substitutions per site and generation, and an assumed generation time of three years – therefore the real values are rough approximations.

**(a)**

| | **P–ST** | **(P, ST)–N** | **N** | **P** | **ST** | **P+ST** | **P+ST+N** |
|---|---|---|---|---|---|---|---|
| | t0 | t1 | q0 | q1 | q2 | q3 | q4 |
| | *Split time (t)* | | *Population size (q)* | | | | |
| Run 1 | 0.456 | 0.538 | 0.0725 | 0.2205 | 0.6690 | 0.0125 | 0.0025 |
| Run 2 | 0.454 | 0.550 | 0.0715 | 0.2195 | 0.6690 | 0.0025 | 0.0125 |
| *Mean* | *0.455* | *0.544* | *0.0720* | *0.2200* | *0.6690* | *0.0075* | *0.0075* |
| | *Split time (years)* | | *Population size (individuals)* | | | | |
| Run 1 | 527369 | 622402 | 6993 | 21270 | 64532 | 1206 | 241 |
| Run 2 | 525980 | 636293 | 6897 | 21173 | 64532 | 241 | 1206 |
| *Mean* | *526675* | *629347* | *6945* | *21221* | *64532* | *723* | *723* |

**(b)**

| **Coalescent direction:** | **N>P** | **P>N** | **N>ST** | **ST>N** | **P>ST** | **ST>P** |
|---|---|---|---|---|---|---|
| Forward direction: | P→N | N→P | ST→N | N→ST | ST→P | P→ST |
| | *Migration rate (m)* | | | | | |
| Run 1 | 0.005 | 0.005 | 4.270 | 0.010 | 2.450 | 0.470 |
| Run 2 | 0.005 | 0.005 | 4.250 | 0.010 | 2.490 | 0.470 |
| *Mean* | *0.005* | *0.005* | *4.260* | *0.010* | *2.470* | *0.470* |
| | *Migration rate (individuals per generation)* | | | | | |
| Run 1 | 0.0002 | 0.0006 | 0.1548 | 0.0033 | 0.2701 | 0.1572 |
| Run 2 | 0.0002 | 0.0005 | 0.1519 | 0.0033 | 0.2733 | 0.1572 |
| *Mean* | *0.0002* | *0.0006* | *0.1534* | *0.0033* | *0.2717* | *0.1572* |

Table S3. Estimates, based on mitochondrial sequences, of the times to the most recent common ancestor (TMRCA). In the second column, estimated times are the most recent common ancestor of the São Tomé grosbeak *Crithagra concolor* and the Príncipe seedeater *C. rufobrunnea*, which are sister species of seedeaters that co-occur in the Gulf of Guinea. The dating in this study is presented for an allopatric speciation scenario, where the São Tomé grosbeak branches off basally to all Príncipe seedeater populations (case 1), and for a scenario in which the São Tomé grosbeak and the São Tomé population of Príncipe seedeater are sister taxa (case 2). The latter could represent sympatric divergence, in which case the TMRCA represents the time since the most recent ancestor, or secondary hybridization, in which case the TMRCA represents the time since last introgression. These results are compared to previous estimates (cases 3–6) from Melo *et al.* (2017), from which estimates are also available for the TMRCA of the entire genus *Crithagra* (third column). All estimates in million years (My) are presented in bold, flanked by the lower and upper bounds of the 95% highest posterior densities, made with Beast. Substitution rates from Lerner *et al.* (2011) except for the 'classic' 2.1% avian substitution rate per My (Weir and Schluter 2008) for a comparison of the cyt *b* dataset. Each case is specified for which loci that were used, followed by the scenario and/or number of taxa included.

| **Case** | **TMRCA (My)** Gulf of Guinea seedeaters | **TMRCA (My)** All *Crithagra* seedeaters |
|---|---|---|
| 1) ATP8&6 – TMRCA 3 populations *C. rufobrunnea* & *C. concolor*: allopatric divergence | 0.59–**0.97**–1.42 | – |
| 2) ATP8&6 – TMRCA *C. concolor* & *C. rufobrunnea* São Tomé: sympatric divergence (or time since last introgression) [a] | 0.41–**0.73**–1.07[b] | – |
| 3) ND2–ND3 & GAPDH–MYO2–ODC: 102 taxa[b] | 0.38–**0.57**–0.77 | 3.11–**3.67**–4.23 |
| 4) ND2: 99 taxa[b] | 0.26–**0.44**–0.65 | 2.30–**2.99**–3.73 |
| 5) Cyt b: 67 taxa[b] | 0.84–**1.31**–1.80 | 3.21–**4.01**–4.86 |
| 6) Cyt b: 67 taxa – 'classic' mtDNA bird clock[b] | 1.06–**1.67**–2.34 | 4.01–**5.11**–6.38 |

[a] This is the estimated divergence time of a sympatric split between *C. concolor* and *C. rufobrunnea* on São Tomé

[b] From Melo *et al.* (2017)

Table S4. Twenty-two genes known to affect bill morphology in birds, and their location in relation to genomic regions that have been preserved from introgression in any direction between the São Tomé grosbeak *Crithagra concolor* and the sympatric population of Príncipe seedeater *C. rufobrunnea* on São Tomé, carrying a clear phylogenetic signal unique to the São Tomé grosbeak (4.6% of the genome; 5 of 24 genes intersecting). For further explanation of the introgressed/preserved genomic regions, see Fig. S4. The genomic location is stated in megabasepairs (Mbp) in relation to the zebra finch *Taeniopygia guttata* (Tgu) genome, assembly taeGut3.2.4.

| **Gene** | **Genomic location (Tgu)** | **Morphological association** | **Reference** |
|---|---|---|---|
| *Bill genes intersecting genomic regions preserved from introgression* | | | |
| HMGA2 | chr1A: 33.5 Mbp | Bill length, depth, width | 1, 2 |
| TGFBR2 | chr2: 60.1 Mbp | Bill length, depth, width | 3, 4, 5 |
| BMP2 | chr3: 26.0 Mbp | Bill depth, width | 6 |
| FGF8 | chr6: 22.1 Mbp | Bill length | 4, 7 |
| COL27A1 | chr17: 5.1 Mbp | Bill length | 8 |
| *Bill genes with no association to preserved regions* | | | |
| ALX1 | chr1A: 41.2 Mbp | Bill pointedness | 9, 10 |
| IGF1 | chr1A: 55.2 Mbp | Bill width | 11 |
| NUP37 | chr1A: 55.3 Mbp | Bill width | 11 |
| WASHC3 | chr1A: 55.4 Mbp | Bill width | 11 |
| PTHRP | chr1A: 72.7 Mbp | Bill length | 12 |
| CTNNB1 | chr2: 64.4 Mbp | Bill length, depth, width | 3, 4, 5 |
| SHH | chr2: 9.0 Mbp | Bill length, depth, width | 4, 7, 13 |
| COL4A5 | chr4A: 12 Mbp | Bill length | 12 |
| DKK3 | chr 5: 1.6 Mbp | Bill length, depth | 3, 5 |
| SOX6 | chr5: 10.8 Mbp | Bill length | 12 |
| CALM1 | chr5: 44.7 Mbp | Bill length, depth | 4, 9, 10, 14 |
| DLK1 | chr5: 50.2 Mbp | Bill size, shape | 2, 10, 15 |
| BMP4 | chr5: 59.4 Mbp | Bill depth, width (length) | 4, 5, 6, 13, 16 |
| IHH | chr7: 10.4 Mbp | Bill length (depth, width) | 5 |
| RALDH2 | chr10: 6.8 Mbp | Bill length, width | 4, 17 |
| RALDH3 | chr10: 18.0 Mbp | Bill length | 17 |
| BMP7 | chr20: 13.0 Mbp | Bill length, depth, width | 4, 6 |

References: 1 – Lamichhaney *et al.* (2016); 2 – Chaves *et al.* (2016); 3 – Mallarino *et al.* (2011); 4 – Knief *et al.* (2012); 5 – Mallarino *et al.* (2012); 6 – Abzhanov *et al.* (2004); 7 – Abzhanov & Tabin (2004); 8 – Cheng *et al.* (2020); 9 – Lamichhaney *et al.* (2015); 10 – Lawson & Petren (2017); 11 – vonHoldt *et al.* (2018); 12 – Bosse *et al.* (2017); 13 – Wu *et al.* (2006); 14 – Abzhanov *et al.* (2006); 15 – Campana *et al.* (2020); 16 – Wu *et al.* (2004); 17 – Song *et al.* (2004).

Table S5. Comparison of microsatellite diversity between the São Tomé grosbeak *Crithagra concolor* (n = 3) and the three Príncipe seedeater *C. rufobrunnea* populations (São Tomé, n = 113; Príncipe, n = 37; Boné, n = 67). The locus Gf10 is not shown as it is Z linked and we only had one male from the São Tomé grosbeak.

| | | | *Crithagra rufobrunnea* | | |
|---|---|---|---|---|---|
| **Locus** | **Het**[a] | *C. concolor* | São Tomé | Príncipe | Boné |
| Ase42 | $H_O$ | 0.67 | 0.81 | 0.41 | 0.14 |
| | $H_E$ | 0.44 | 0.74 | 0.33 | 0.18 |
| Ase43 | $H_O$ | 1.00 | 0.69 | 0.81 | 0.58 |
| | $H_E$ | 0.63 | 0.83 | 0.74 | 0.70 |
| Ase48 | $H_O$ | 0.33 | 0.79 | 0.73 | 0.70 |
| | $H_E$ | 0.61 | 0.88 | 0.69 | 0.73 |
| Cuμ104 | $H_O$ | 0.67 | 0.72 | 0.11 | 0.00 |
| | $H_E$ | 0.61 | 0.71 | 0.16 | 0.00 |
| Cuμ128 | $H_O$ | 0.67 | 0.86 | 0.69 | 0.83 |
| | $H_E$ | 0.61 | 0.88 | 0.80 | 0.72 |
| Gf08 | $H_O$ | 0.33 | 0.17 | 0.03 | 0.00 |
| | $H_E$ | 0.28 | 0.23 | 0.03 | 0.00 |
| LOX1 | $H_O$ | 1.00 | 0.96 | 0.92 | 0.27 |
| | $H_E$ | 0.75 | 0.95 | 0.89 | 0.44 |
| LOX3 | $H_O$ | 1.00 | 0.99 | 0.84 | 0.84 |
| | $H_E$ | 0.83 | 0.98 | 0.93 | 0.82 |
| LOC7 | $H_O$ | 1.00 | 0.98 | 0.97 | 0.66 |
| | $H_E$ | 0.83 | 0.98 | 0.97 | 0.92 |
| LOX8 | $H_O$ | 1.00 | 0.92 | 0.58 | 0.42 |
| | $H_E$ | 0.72 | 0.97 | 0.91 | 0.62 |
| Lswμ14 | $H_O$ | 0.00 | 0.33 | 0.22 | 0.16 |
| | $H_E$ | 0.00 | 0.52 | 0.31 | 0.26 |
| Lswμ18 | $H_O$ | 1.00 | 0.51 | 0.73 | 0.02 |
| | $H_E$ | 0.78 | 0.69 | 0.85 | 0.02 |
| Pdoμ4 | $H_O$ | 1.00 | 0.77 | 0.92 | 0.76 |
| | $H_E$ | 0.83 | 0.82 | 0.95 | 0.82 |
| WBSW7 | $H_O$ | 0.67 | 0.57 | 0.00 | 0.00 |
| | $H_E$ | 0.44 | 0.55 | 0.00 | 0.00 |

[a] Heterozygosity: observed ($H_O$) and expected ($H_E$).

Table S6. Sample details for birds sequenced in this study. As identifiers, the main sample number (FIELD) and ring number are presented. Date is given as YYYY-MM-DD. Abbreviations: altitude (Alt.); male (M); female (F).

| FIELD | Ring | Taxon | Sex | Date | Country | Province | Locality | Coordi-nates | Alt. (m) | Collector |
|---|---|---|---|---|---|---|---|---|---|---|
| P1.141 | FA33535 | *Crithagra r. rufobrunnea* | M | 2002-12-13 | São Tomé & Príncipe | Príncipe | Ribeira Porco | 0132N 0722E | 128 | M. Melo |
| P1.142 | FA33536 | *Crithagra r. rufobrunnea* | M | 2002-12-13 | São Tomé & Príncipe | Príncipe | Ribeira Porco | 0132N 0722E | 128 | M. Melo |
| P1.266 | FA33738 | *Crithagra r. rufobrunnea* | M | 2003-04-03 | São Tomé & Príncipe | Príncipe | Ribeira Porco | 0132N 0722E | 128 | M. Melo |
| P1.267 | FA33739 | *Crithagra r. rufobrunnea* | M | 2003-04-03 | São Tomé & Príncipe | Príncipe | Ribeira Porco | 0132N 0722E | 128 | M. Melo |
| P2.021 | FA33770 | *Crithagra r. rufobrunnea* | F | 2003-12-14 | São Tomé & Príncipe | Príncipe | Praia Cará | 0133N 0721E | 27 | M. Melo |
| P3.026 | FA33889 | *Crithagra r. rufobrunnea* | M | 2004-12-16 | São Tomé & Príncipe | Príncipe | Praia Cará | 0133N 0721E | 27 | M. Melo |
| P3.027 | FA33890 | *Crithagra r. rufobrunnea* | F | 2004-12-16 | São Tomé & Príncipe | Príncipe | Praia Cará | 0133N 0721E | 27 | M. Melo |
| P3.030 | FA33891 | *Crithagra r. rufobrunnea* | M | 2004-12-17 | São Tomé & Príncipe | Príncipe | Praia Cará | 0133N 0721E | 27 | M. Melo |
| P3.031 | FA33892 | *Crithagra r. rufobrunnea* | M | 2004-12-17 | São Tomé & Príncipe | Príncipe | Praia Cará | 0133N 0721E | 27 | M. Melo |
| P3.032 | FA33893 | *Crithagra r. rufobrunnea* | M | 2004-12-17 | São Tomé & Príncipe | Príncipe | Praia Cará | 0133N 0721E | 27 | M. Melo |
| P3.041 | FA33903 | *Crithagra r. rufobrunnea* | F | 2004-12-17 | São Tomé & Príncipe | Príncipe | Praia Cará | 0133N 0721E | 27 | M. Melo |
| P3.045 | FA33894 | *Crithagra r. rufobrunnea* | F | 2004-12-17 | São Tomé & Príncipe | Príncipe | Praia Cará | 0133N 0721E | 27 | M. Melo |
| P3.055 | FA33910 | *Crithagra r. rufobrunnea* | F | 2004-12-19 | São Tomé & Príncipe | Príncipe | Praia Cará | 0133N 0721E | 27 | M. Melo |
| P3.057 | FA33911 | *Crithagra r. rufobrunnea* | F | 2004-12-19 | São Tomé & Príncipe | Príncipe | Praia Cará | 0133N 0721E | 27 | M. Melo |
| P5.030 | FA92819 | *Crithagra r. rufobrunnea* | | 2011-07-31 | São Tomé & Príncipe | Príncipe | Praia Cará | 0133N 0721E | 27 | M. Melo, M. Stervander |
| P5.031 | FA92820 | *Crithagra r. rufobrunnea* | | 2011-08-01 | São Tomé & Príncipe | Príncipe | Praia Cará | 0133N 0721E | 27 | M. Melo, M. Stervander |
| P5.034 | FA92821 | *Crithagra r. rufobrunnea* | | 2011-08-01 | São Tomé & Príncipe | Príncipe | Praia Cará | 0133N 0721E | 27 | M. Melo, M. Stervander |
| P5.035 | FA92822 | *Crithagra r. rufobrunnea* | | 2011-08-01 | São Tomé & Príncipe | Príncipe | Praia Cará | 0133N 0721E | 27 | M. Melo, M. Stervander |
| P5.038 | FA92825 | *Crithagra r. rufobrunnea* | | 2011-08-01 | São Tomé & Príncipe | Príncipe | Praia Cará | 0133N 0721E | 27 | M. Melo, M. Stervander |
| P5.042 | FA92826 | *Crithagra r. rufobrunnea* | | 2011-08-01 | São Tomé & Príncipe | Príncipe | Praia Cará | 0133N 0721E | 27 | M. Melo, M. Stervander |
| P5.045 | FA92827 | *Crithagra r. rufobrunnea* | | 2011-08-02 | São Tomé & Príncipe | Príncipe | Praia Cará | 0133N 0721E | 27 | M. Melo, M. Stervander |
| P3.058 = P5.047 | FA33912 | *Crithagra r. rufobrunnea* | | 2011-08-02 | São Tomé & Príncipe | Príncipe | Praia Cará | 0133N 0721E | 27 | M. Melo, M. Stervander |
| P1.231 | FA33538 | *Crithagra r. fradei* | M | 2002-12-20 | São Tomé & Príncipe | Príncipe | Boné de Jóquei | 0130N 0725E | 5 | M. Melo |
| P1.232 | FA33539 | *Crithagra r. fradei* | M | 2002-12-20 | São Tomé & Príncipe | Príncipe | Boné de Jóquei | 0130N 0725E | 5 | M. Melo |
| P1.233 | FA33540 | *Crithagra r. fradei* | M | 2002-12-21 | São Tomé & Príncipe | Príncipe | Boné de Jóquei | 0130N 0725E | 5 | M. Melo |
| P1.234 | FA33541 | *Crithagra r. fradei* | F | 2002-12-21 | São Tomé & Príncipe | Príncipe | Boné de Jóquei | 0130N 0725E | 5 | M. Melo |
| P1.235 | FA33542 | *Crithagra r. fradei* | M | 2002-12-21 | São Tomé & Príncipe | Príncipe | Boné de Jóquei | 0130N 0725E | 5 | M. Melo |
| P1.236 | FA33543 | *Crithagra r. fradei* | F | 2002-12-21 | São Tomé & Príncipe | Príncipe | Boné de Jóquei | 0130N 0725E | 5 | M. Melo |
| P1.239 | FA33546 | *Crithagra r. fradei* | F | 2002-12-21 | São Tomé & Príncipe | Príncipe | Boné de Jóquei | 0130N 0725E | 5 | M. Melo |
| P1.243 | FA33551 | *Crithagra r. fradei* | M | 2002-12-21 | São Tomé & Príncipe | Príncipe | Boné de Jóquei | 0130N 0725E | 5 | M. Melo |
| P1.249 | FA33556 | *Crithagra r. fradei* | M | 2002-12-21 | São Tomé & Príncipe | Príncipe | Boné de Jóquei | 0130N 0725E | 5 | M. Melo |
| P1.250 | FA33557 | *Crithagra r. fradei* | F | 2002-12-21 | São Tomé & Príncipe | Príncipe | Boné de Jóquei | 0130N 0725E | 5 | M. Melo |
| P1.259 | FA33566 | *Crithagra r. fradei* | F | 2002-12-21 | São Tomé & Príncipe | Príncipe | Boné de Jóquei | 0130N 0725E | 5 | M. Melo |
| P1.263 | dead | *Crithagra r. fradei* | F | 2002-12-21 | São Tomé & Príncipe | Príncipe | Boné de Jóquei | 0130N 0725E | 5 | M. Melo |
| P2.002 | FA33758 | *Crithagra r. fradei* | F | 2003-12-12 | São Tomé & Príncipe | Príncipe | Boné de Jóquei | 0130N 0725E | 5 | M. Melo |
| P3.001 | FA33863 | *Crithagra r. fradei* | F | 2004-12-15 | São Tomé & Príncipe | Príncipe | Boné de Jóquei | 0130N 0725E | 5 | M. Melo |
| P3.004 | FA33866 | *Crithagra r. fradei* | F | 2004-12-15 | São Tomé & Príncipe | Príncipe | Boné de Jóquei | 0130N 0725E | 5 | M. Melo |
| P5.005 | FA62400 | *Crithagra r. fradei* | | 2011-07-30 | São Tomé & Príncipe | Príncipe | Boné de Jóquei | 0130N 0725E | 5 | M. Melo, M. Stervander |
| P5.006 | FA62401 | *Crithagra r. fradei* | | 2011-07-30 | São Tomé & Príncipe | Príncipe | Boné de Jóquei | 0130N 0725E | 5 | M. Melo, M. Stervander |
| P5.011 | FA62406 | *Crithagra r. fradei* | | 2011-07-30 | São Tomé & Príncipe | Príncipe | Boné de Jóquei | 0130N 0725E | 5 | M. Melo, M. Stervander |
| P5.016 | FA62411 | *Crithagra r. fradei* | | 2011-07-31 | São Tomé & Príncipe | Príncipe | Boné de Jóquei | 0130N 0725E | 5 | M. Melo, M. Stervander |
| P5.010 | FA62405 | *Crithagra r. fradei* | | 2011-07-30 | São Tomé & Príncipe | Príncipe | Boné de Jóquei | 0130N 0725E | 5 | M. Melo, M. Stervander |

| | | | | | | | | | | |
|---|---|---|---|---|---|---|---|---|---|---|
| P5.014 | FA62409 | *Crithagra r. fradei* | | 2011-07-30 | São Tomé & Príncipe | Príncipe | Boné de Jóquei | 0130N 0725E | 5 | M. Melo, M. Stervander |
| P5.015 | FA62410 | *Crithagra r. fradei* | | 2011-07-31 | São Tomé & Príncipe | Príncipe | Boné de Jóquei | 0130N 0725E | 5 | M. Melo, M. Stervander |
| P5.028 | FA62418 | *Crithagra r. fradei* | | 2011-07-31 | São Tomé & Príncipe | Príncipe | Boné de Jóquei | 0130N 0725E | 5 | M. Melo, M. Stervander |
| ST1.044 | FA33506 | *Crithagra r. thomensis* | F | 2002-10-31 | São Tomé & Príncipe | São Tomé | Monte Café | 0018N 0638E | 723 | M. Melo |
| ST1.100 | FA33509 | *Crithagra r. thomensis* | F | 2002-11-14 | São Tomé & Príncipe | São Tomé | Pico Calvário | 0016N 0634E | 1587 | M. Melo |
| ST1.125 = | FA33516 | *Crithagra r. thomensis* | M | 2002-11-20 | São Tomé & Príncipe | São Tomé | Alto Douro | 0012N 0641E | 150 | M. Melo |
| ST5.027 (recapture) | | *Crithagra r. thomensis* | | 2011-07-13 | São Tomé & Príncipe | São Tomé | Alto Douro | 0012N 0641E | 150 | M. Melo, M. Stervander |
| ST1.146 | FA33524 | *Crithagra r. thomensis* | F | 2002-11-20 | São Tomé & Príncipe | São Tomé | Alto Douro | 0012N 0641E | 150 | M. Melo |
| ST1.204 | FA33532 | *Crithagra r. thomensis* | F | 2002-11-27 | São Tomé & Príncipe | São Tomé | Bom Sucesso | 0017N 0636E | 1158 | M. Melo |
| ST1.225 | FA33576 | *Crithagra r. thomensis* | M | 2002-12-29 | São Tomé & Príncipe | São Tomé | Quija | 0009N 0633E | 658 | M. Melo |
| ST1.242 | FA33604 | *Crithagra r. thomensis* | M | 2003-01-07 | São Tomé & Príncipe | São Tomé | Lagoa Amélia | 0016N 0635E | 1498 | M. Melo |
| ST1.312 | FA33582 | *Crithagra r. thomensis* | F | 2003-01-24 | São Tomé & Príncipe | São Tomé | Quija | 0009N 0633E | 658 | M. Melo |
| ST2.108 | FA33788 | *Crithagra r. thomensis* | F | 2003-12-29 | São Tomé & Príncipe | São Tomé | Lagoa Amélia | 0016N 0635E | 1498 | M. Melo |
| ST2.157 | FA33797 | *Crithagra r. thomensis* | F | 2004-01-04 | São Tomé & Príncipe | São Tomé | Umbumgu | 0009N 0633E | 300 | M. Melo |
| ST2.256 | FA33857 | *Crithagra r. thomensis* | F | 2004-03-07 | São Tomé & Príncipe | São Tomé | Alto Douro | 0012N 0641E | 150 | M. Melo |
| ST3.012 | FA33917 | *Crithagra r. thomensis* | F | 2005-01-05 | São Tomé & Príncipe | São Tomé | Contador dam | 0019N 0632E | 600 | M. Melo |
| ST3.015 | FA33918 | *Crithagra r. thomensis* | M | 2005-01-05 | São Tomé & Príncipe | São Tomé | Contador dam | 0019N 0632E | 600 | M. Melo |
| ST3.295 | FA62325 | *Crithagra r. thomensis* | M | 2005-02-10 | São Tomé & Príncipe | São Tomé | São Miguel | 0010N 0630E | 400 | M. Melo |
| ST3.342 | FA62340 | *Crithagra r. thomensis* | F | 2005-02-12 | São Tomé & Príncipe | São Tomé | São Miguel | 0010N 0630E | 400 | M. Melo |
| ST5.023 | FA62383 | *Crithagra r. thomensis* | | 2011-07-12 | São Tomé & Príncipe | São Tomé | Alto Douro | 0012N 0641E | 150 | M. Melo, M. Stervander |
| ST5.026 | FA62385 | *Crithagra r. thomensis* | | 2011-07-13 | São Tomé & Príncipe | São Tomé | Alto Douro | 0012N 0641E | 150 | M. Melo, M. Stervander |
| ST5.097 | FH58937 | *Crithagra r. thomensis* | | 2011-08-16 | São Tomé & Príncipe | São Tomé | Calvário Bend | 0016N 0635E | 1363 | M. Melo, M. Stervander |
| ST5.098 | FH58938 | *Crithagra r. thomensis* | | 2011-08-16 | São Tomé & Príncipe | São Tomé | Calvário Bend | 0016N 0635E | 1363 | M. Melo, M. Stervander |
| ST5.100 | FH58940 | *Crithagra r. thomensis* | | 2011-08-16 | São Tomé & Príncipe | São Tomé | Calvário Bend | 0016N 0635E | 1363 | M. Melo, M. Stervander |
| ST5.123 | FH58956 | *Crithagra r. thomensis* | | 2011-08-17 | São Tomé & Príncipe | São Tomé | Calvário Bend | 0016N 0635E | 1363 | M. Melo, M. Stervander |
| ST5.125 | FH58958 | *Crithagra r. thomensis* | | 2011-08-17 | São Tomé & Príncipe | São Tomé | Calvário Bend | 0016N 0635E | 1363 | M. Melo, M. Stervander |
| ST1.362 | FA33592 | *Crithagra concolor* | F | 2003-01-27 | São Tomé & Príncipe | São Tomé | Umbumgu | 0009N 0633E | 300 | M. Melo |
| ST3.318 | BE06749 | *Crithagra concolor* | M | 2005-02-11 | São Tomé & Príncipe | São Tomé | São Miguel | 0010N 0630E | 400 | M. Melo |
| ST3.319 | BE06800 | *Crithagra concolor* | M | 2005-02-11 | São Tomé & Príncipe | São Tomé | São Miguel | 0010N 0630E | 400 | M. Melo |
| ST5.039 | 4A38193 | *Crithagra concolor* | F | 2011-07-26 | São Tomé & Príncipe | São Tomé | São Miguel | 0010N 0630E | 400 | M. Melo |
| C084 | FA33599 | *Crithagra burtoni* | M | 2003-02-16 | Cameroon | Limbe | Mann's Spring | 0408N 0907E | 2282 | M. Melo |
| C119 | FA33703 | *Crithagra burtoni* | | 2003-02-17 | Cameroon | Limbe | Mann’s Spring | 0408N 0907E | 2282 | M. Melo |
| C133 | FA33705 | *Crithagra burtoni* | M | 2003-02-17 | Cameroon | Limbe | Mann's Spring | 0408N 0907E | 2282 | M. Melo |
| C159 | FA33712 | *Crithagra burtoni* | M | 2003-02-17 | Cameroon | Limbe | Mann's Spring | 0408N 0907E | 2282 | M. Melo |
| Z108 | GA59089 | *Crithagra flaviventris* | M | 2002-09-12 | South Africa | NC | Vanzyls-rus | | | Claire Spottiswoode |
| Z111 | | *Crithagra flaviventris* | M | 2003-12-27 | South Africa | WC | Dwaal-hoek | | | Claire Spottiswoode |

Table S7. PCR primer details and evaluation of amplification success in eight passerine bird species, including three seedeaters (shadowed grey), ordered according to position in the zebra finch *Taeniopygia guttata* genome. Unless referenced, primers were designed for this study. Zebra finch (Tgu) loci are annotated according to the taeGut3.2.4 assembly (https://sep2019.archive.ensembl.org/Taeniopygia_guttata/Info/Index): coding sequence (cds); untranslated region (utr); downstream (ds). Position in the zebra finch genome is given by chromosome and basepair (bp) for starting nucleotide. Length in zebra finch refers to intron length for primers amplifying introns; otherwise to total product length (without primers). Amplification success is scored as good amplification (+), no amplification (–), or multiple/unspecific amplification (m).

| | | | | | | | | Amplification success† | | | | | | | |
|---|---|---|---|---|---|---|---|---|---|---|---|---|---|---|---|
| **Marker** *Tgu locus: segment* | **Gene position Tgu (chromosome: bp start)** | | **Length (bp) *T. guttata*** | **Reference*** | **Primer sequence (5'–3')** | | **Annealing temperature (°C)** | *A. arundinaceus* | *C. caeruleus* | *C. corone* | *T. guttata* | *C. flaviventris* | *C. burtoni* | *C. rufobrunnea* | *N. acunhae* |
| ENSGALT00000026187 | 1: | 6,882,519 | 795 | 1 | F: | GGTGGGAATGCAGTAGTAGA | 63–53 | + | + | + | + | + | + | + | + |
| *ATP6AP2: intron 7* | | | | | R: | CATGTTCCAGAGGTTGTAGG | | | | | | | | | |
| ENSGALT00000026698 | 1: | 16,280,498 | 513 | 1 | F: | CAGAGGATGCGGAAGATGG | 63–53 | + | m | + | + | + | + | + | + |
| *RBBP7: intron 12* | | | | | R: | TGATACAGAACAGATGACCC | | | | | | | | | |
| UBE3A | 1: | 31,837,212 | 471 | 8 | F: | AGAAACTACAGAATATGATGGTGGC | 59.9 | + | + | + | m | + | + | + | + |
| *UBE3A: intron 12* | | | | | R: | CTGTCTGTGCCTGTTGTAAACTG | | | | | | | | | |
| NBEA | 1: | 52,371,012 | 640 | | F: | TTTTAACCGGTGGCATTATGA | 60.2 | + | + | + | + | + | + | + | + |
| *NBEA: exon 56 (cds) + downstream* | | | | | R: | CTRTTTGCKCCCAAATTC | | | | | | | | | |
| ALG11 | 1: | 54,992,341 | 719 | | F: | GARGCRGCCACTGCTGGT | 60.0 | m | m | m | m | + | + | + | m |
| *ALG11: intron 1* | | | | | R: | TTGAATCTTCTGAAAGCACCTTC | | | | | | | | | |
| ENSGALT00000027331 | 1: | 71,230,345 | 670 | 1 | F: | CCTAGCTAAATATGTTCTGGC | 60–50 | + | + | + | + | m | + | + | + |
| *RBM26: intron 2* | | | | | R: | TAGGCTTCCTGATGATGGCT | | | | | | | | | |
| GADPH | 1: | 90,242,467 | 274 | 2 | F: | ACCTTTCATGCGGGTGCTGGCATTGC | 64.0 | + | + | + | + | + | + | + | + |
| *GAPDH: intron 10* | | | | | R: | CATCAAGTCCACAACACGGTTGCTGTA | | | | | | | | | |
| APP | 1: | 112,697,462 | 724 | | F: | GCTTCACTGCCCATTGGT | 58.7 | + | + | + | + | + | + | + | + |
| *APP: exon 17 (utr) + downstream* | | | | | R: | TAAGCAATGGYTCTGTACAATCA | | | | | | | | | |
| ENSGALT00000018851 | 1A: | 46,566,037 | 772 | 1 | F: | CAGTTTCAAGCAGTTCTTAG | 63–53 | + | + | + | + | + | + | + | + |
| *ACTR6: intron 3* | | | | | R: | GATACAACACAGCTCAGATG | | | | | | | | | |
| Tgu1A-49.14MbpEx | 1A: | 49,140,734 | 757 | | F: | CTATAGAGAACCRYGTGGACTT | 60.0 | + | + | + | + | + | + | + | + |
| *Not_annotated: exon (not annotated)* | | | | | R: | AYTGGAAGAGAATCAGTTCCCTC | | | | | | | | | |

| | | | | | | | | | | | | | | | |
|---|---|---|---|---|---|---|---|---|---|---|---|---|---|---|---|
| Myo2 | 1A: | 51,873,668 | 699 | 3 | F: | GCCACCAAGCACAAGATCCC | 58.0 | + | + | + | + | + | m | + | + |
| *MB: intron 2* | | | | 4 | R: | GCAAGGACCTTGATAATGACTT | | | | | | | | | |
| LDH-B | 1A: | 65,040,952 | 488 | 5 | F: | GGAAGACAAACTAAAAGGAGAAATGATGGA | 0.0 | + | + | m | + | + | + | + | + |
| *LDHB: intron 3* | | | | | R: | TTCCTCTGAAGCAGGTTGAGACGACTCTC | | | | | | | | | |
| ZEB1_ds | 2: | 15,888,805 | 688 | | F: | AAAAACACTCAGCCATCAGAGAA | 59.7 | + | + | + | + | + | + | + | + |
| *ZEB1_ds: downstream (exon, utr?)* | | | | | R: | TCTCTTAAYAAAAYTGCAGAGCA | | | | | | | | | |
| LRPP4/RP40 | 2: | 63,479,351 | 221 | 2, 5 | F: | GGGCCTGATGTGGTGGATGCTGGC | 67.0 | + | + | + | + | + | + | + | + |
| *RPSA: intron 7* | | | | | R: | GCTTTCTCAGCAGCAGCCTGCTC | | | | | | | | | |
| ENSGALT00000020352 | 2: | 76,673,689 | 1014 | 1 | F: | TCCTTATCAAGCAGCAGTAG | 60–50 | + | + | + | + | + | m | + | + |
| *HUS1: intron 4* | | | | | R: | CTTCATTGTTTTCAGCACTG | | | | | | | | | |
| PDCD6 | 2: | 94,138,715 | 678 | 8 | F: | TGGAARTACATCACAGAYTGGC | 60.0 | + | + | + | + | + | + | + | + |
| *PDCD6: intron 4* | | | | | R: | ACACAGCACTGAATAAARTCATCAA | | | | | | | | | |
| PENK | 2: | 117,404,415 | 759 | | F: | TGCTTGCMAAGAARTACGG | 60.0 | m | + | m | + | + | + | + | + |
| *PENK: exon 2 (cds+utr)* | | | | | R: | ACAGTWAAGCAAYTCTACTGTRGATG | | | | | | | | | |
| RPL7 | 2: | 124,033,847 | 634 | 8 | F: | TACCATAAGGAGTACAGRCACATG | 60.2 | – | + | + | + | + | + | + | + |
| *RPL7: intron 3* | | | | | R: | TAGGGYTCAACAATCCGCA | | | | | | | | | |
| ENSGALT00000025969[a] | 2: | 142,002,162 | 889 | 1 | F: | GGCAACCCTCATCACTTCT | 65–55 | + | – | + | + | + | + | + | + |
| *UTP23: intron 2* | | | | | R: | CTAGCACCATAGTGTTCTGA | | | | | | | | | |
| ENSGALT00000026472 | 2: | 143,445,412 | 874 | 1 | F: | TCTGAACCACTTACTGGCAG | 63–53 | + | + | + | m | + | + | + | + |
| *COL14A1: intron 14* | | | | | R: | CAAAGGAATCCATGCCAATC | | | | | | | | | |
| TGFb2 | 3: | 10,256,872 | 575 | 2 | F: | GAAGCGTGCTCTAGATGCTG | 65.0 | + | + | + | + | + | + | + | + |
| *TGFB2: intron 6* | | | | | R: | AGGCAGCAATTATCCTGCAC | | | | | | | | | |
| ENSGALT00000016264 | 3: | 18,103,287 | 680 | 1 | F: | GCATTGACCTCAAAGAAGGC | 63–53 | + | + | + | m | + | + | + | + |
| *CRIPT: intron 3* | | | | | R: | TTTATAGGCACATCCTTGAC | | | | | | | | | |
| SNAP25 | 3: | 24595961 | 821 | | F: | TTGGAATGCTAAACGTGCTG | 60.1 | + | + | + | + | + | + | + | + |
| *SNAP25: exon 8 (utr)* | | | | | R: | ACAAAATGTCAATCACTTCACAAACT | | | | | | | | | |
| USP34 | 3: | 26,405,624 | 886 | | F: | RTGTGCACGTAGATGGGAATC | 59.9 | + | + | – | + | + | + | + | + |
| *USP34: intron 816* | | | | | R: | TCAGTAAACTTYTCACAGYTGATTG | | | | | | | | | |
| PCMT1 | 3: | 57,196,709 | 1585 | | F: | GAGATGGAMGAATGGGAT | 60.1 | m | + | m | + | + | – | + | + |
| *PCMT1: intron 6* | | | | | R: | CWGCAGGRCCAACTGGTA | | | | | | | | | |
| RNF146 | 3: | 61,489,959 | 781 | | F: | AGGCAATRGAGAATACGCTTGG | 59.9 | + | + | + | + | + | + | + | + |
| *RNF146: exon 1 (cds)* | | | | | R: | TGCAATGCCAATGYGGAC | | | | | | | | | |
| AKIRIN2 | 3: | 78,683,627 | 699 | | F: | GGAGTACAAACGCATGCAGA | 60.0 | + | + | + | + | + | + | + | + |
| *AKIRIN2: intron 1* | | | | | R: | ACTCTTCACGGATTTTCTCTTCAC | | | | | | | | | |

| | | | | | | | | | | | | | | |
|---|---|---|---|---|---|---|---|---|---|---|---|---|---|---|
| ODC | 3: | 99,482,257 | 527 | 2 | F: | GACTCCAAAGCAGTTTGTCGTCTCAGTGT | 60.0 | + | + | + | + | + | + | + | + |
| *ODC1: intron 4/exon 5/intron 5* | | | | | R: | TCTTCAGAGCCAGGGAAGCCACCACCAAT | | | | | | | | | |
| PCDH18 | 4: | 124,992 | 948 | | F: | TCRCTGAACAGCCTGGTGAC | 59.7 | + | + | + | + | + | + | + | + |
| *PCDH18: intron 1* | | | | | R: | AATAYTTGTTTCCTCKAAAACTTGGC | | | | | | | | | |
| ANKRD17[b] | 4: | 1,459,256 | 890 | | F: | YAGCAYGTTGATCCCACTC | 60.3 | + | + | + | + | m | m | m | + |
| *ANKRD17: intron 43* | | | | | R: | CTCATGCTGCTGACCCATC | | | | | | | | | |
| Tgu4-30.05MbpEx | 4: | 30,052,158 | 698 | | F: | TGCATGTGTAGAGAAGGCCA | 60.4 | + | + | + | + | + | + | + | + |
| *Not_annotated: exon (not annotated)* | | | | | R: | CCAGCTTCATACAAACTGGATGT | | | | | | | | | |
| MPLPR | 4A: | 19,585,309 | 211 | 2 | F: | TACATCTACTTTAACACCTGGACCACCTG | 65.0 | + | + | + | + | + | + | + | + |
| *PLP1: intron 8* | | | | | R: | TTGCAGATGGAGAGCAGGTTGGAGCC | | | | | | | | | |
| ENSGALT00000025924 | 4_random: | 4,029,445 | 652 | 1 | F: | TTCATCCAGGACCAGATCGC | 60–50 | m | + | + | + | m | + | + | + |
| *C2orf7 : intron 3* | | | | | R: | GTGCTGTCCTGGATCATCTC | | | | | | | | | |
| EIF4G2 | 5: | 8,245,723 | 778 | | F: | ACCATAMCCAGAATCAGGGAC | 59.9 | + | + | – | + | + | + | + | + |
| *EIF4G2: intron 13* | | | | | R: | GGTTGAGCACTGGGAGGA | | | | | | | | | |
| CHP1 | 5: | 23,401,995 | 1235 | | F: | GACTTCCAGCGGATYCCA | 59.9 | + | + | – | + | + | + | + | + |
| *CHP: intron 3* | | | | | R: | TCTGGTCCATTCTGGTCTTTG | | | | | | | | | |
| NOVA1_ds | 5: | 32,518,453 | 669 | | F: | ACCAGGAGCTTTTCTTTCAGC | 60.1 | + | + | + | + | + | + | + | + |
| *NOVA1_ds: downstream (exon, utr?)* | | | | | R: | GCATATWGTAATTTGCATGACCCA | | | | | | | | | |
| USP47 | 5_random: | 1,647,991 | 667 | | F: | AAAAGAAAGCAGCAGACTCCAG | 60.1 | + | m | m | + | m | m | m | –/m |
| *USP47: exon 28 (cds) + downstream* | | | | | R: | ACCTTACAACAGCACTTCTGCA | | | | | | | | | |
| CDC2 | 6: | 5,597,471 | 538 | | F: | CCTCAGAAYCTYCTAATAGATGACA | 60.3 | + | + | + | + | + | + | + | + |
| *CDC2: intron 4* | | | | | R: | TAACGAGCAGATCCYAGCA | | | | | | | | | |
| OGDHL | 6: | 8,571,613 | 692 | 8 | F: | GCAGTTAGAATGTCTTTCCCTCA | 59.8 | + | + | + | + | + | + | + | + |
| *OGDHL: intron 21* | | | | | R: | GGTGTCAGGAGGARCACAA | | | | | | | | | |
| TM9SF3_ds | 6: | 15,585,498 | 727 | | F: | GTGAAGAGGACAAACRGTTTRTTTA | 60.2 | + | + | + | + | + | + | + | + |
| *TM9SF3_ds: downstream (exon, utr?)* | | | | | R: | CAATGGCAGGCARGATGT | | | | | | | | | |
| ENSGALT00000015724 | 6: | 32,012,042 | 693 | 1 | F: | AAATGTATTGAATGGATGGG | 60–50 | + | + | + | + | + | + | + | + |
| *ACADSB: intron 9* | | | | | R: | CAGCTGGATATTTGAAGTTC | | | | | | | | | |
| ARPC2 | 7: | 10,063,664 | 419 | | F: | TGAAGCGCAACTGCTTTG | 59.9 | + | + | – | + | + | + | + | + |
| *ARPC2: intron 5* | | | | | R: | GCATAAACACCTTTCCGATCA | | | | | | | | | |
| WDR12 | 7: | 21,530,614 | 530 | 8 | F: | TGACTTCTCACACAGGCTGG | 60.0 | + | + | + | + | + | + | + | + |
| *WDR12: intron 9* | | | | | R: | CATGAGCAGCCAAGTCATACA | | | | | | | | | |
| DPP10 | 7: | 31,509,183 | 659 | | F: | CCAAGGGACAAAACCTTTACCT | 60.2 | + | + | + | + | + | + | + | + |
| *DPP10: presumed 3'-utr* | | | | | R: | CTAAAATTAGAGTGYAACCAGCAA | | | | | | | | | |

| | | | | | | | | | | | | | | | |
|---|---|---|---|---|---|---|---|---|---|---|---|---|---|---|---|
| WDR47 | 8: | 5,806,868 | 572 | | F: | NGGCAGTTGATTCTTGATGG | 59.7 | + | + | + | + | + | + | + | + |
| *WDR47: intron 2* | | | | | R: | TGCATTGTTYACACATAAGGC | | | | | | | | | |
| CNN3 | 8: | 9,792,282 | 726 | | F: | AAGAGATTACCATGGTCAGTACAGTG | 59.9 | + | + | + | + | + | + | + | + |
| *CNN3: presumed 3'-utr* | | | | | R: | AATACAMTGGGATCAGTTTSAGAAG | | | | | | | | | |
| BTF3L4 | 8: | 21,805,788 | 855 | 8 | F: | GGAYAGTAAAGCACCAAAATCTGAA | 60.1 | + | + | + | + | + | + | + | + |
| *BTF3L4: intron 6* | | | | | R: | GTCCATGCCAGCTTCCAG | | | | | | | | | |
| WDR33 | 9: | 4,122,095 | 632 | | F: | TGCTGAACAAYCCTATGAATGC | 60.0 | + | + | m | + | + | + | + | + |
| *WDR33: intron 4* | | | | | R: | AAATTGAAAGTCAGTCCATTCCA | | | | | | | | | |
| GUCY1B2 | 9: | 9,576,523 | 1820 | | F: | TCAATCAAACRGARGAAGAAGAAAGA | 59.9 | + | + | – | + | + | + | + | + |
| *GUCY1B2: intron 6/exon 7/intron 7* | | | | | R: | TGCAAAATGTTTTTGTGTCAATC | | | | | | | | | |
| SSR3 | 9: | 26,218,883 | 440 | | F: | GTTCAGTCTGCMGTSCTGTA | 60.4 | + | + | + | + | m | m | m | + |
| *SSR3: intron 2* | | | | | R: | CTTCTCCTTSCGRGACATC | | | | | | | | | |
| ENSGALT00000006419 | 10: | 5,882,161 | 516 | 1 | F: | AGCAATGGGCTTTGCAGTAG | 63–53 | + | + | – | + | + | + | + | + |
| *TM2D3: intron 2* | | | | | R: | CTGGCACGTCATGTTAATTG | | | | | | | | | |
| DNAJA2 | 11: | 13,887,105 | 460 | 8 | F: | ACAGTATCGCAATCCTTTTGAGA | 60.4 | + | + | + | + | + | + | + | + |
| *DNAJA2: intron 3* | | | | | R: | TCTGGCCACCACCTGAAC | | | | | | | | | |
| ENSGALT00000011836[c] | 12: | 14,512,888 | 573 | 1 | F: | GAAAAGATCTTGGGGGAGAG | 63–53 | + | + | m | m | + | + | + | + |
| *PSMD6: intron part 5* | | | | | R: | CTTGCGGAATGCAGTCAGAG | | | | | | | | | |
| Rho1 | 12: | 21,214,088 | 926 | 2 | F: | TGCTACATCGAGGGCTTCTT | 56.0 | + | m | – | + | + | + | + | + |
| *RHO: intron 1* | | | | | R: | GCAGTGACCAGAGAGCGATT | | | | | | | | | |
| PPP2CA | 13: | 10,139,310 | 653 | | F: | AGAGGTGAACCGCATGTTACTC | 60.0 | m | + | + | + | + | + | + | + |
| *PPP2CA: exon 7 (mainly utr)* | | | | | R: | AAAAACACTGCAGCTCTGTCC | | | | | | | | | |
| MATR3 | 13_random: | 601,295 | 586 | | F: | CAAGATTGAGGAGCCAGAGC | 60.1 | + | + | m | m | + | + | + | + |
| *MATR3: intron 12* | | | | | R: | AACTTTTGATAATGAGGAAGGCTG | | | | | | | | | |
| A2BP1 | 14: | 4,502,086 | 658 | | F: | GCAGTAGTGCATCATTTTAGCAAC | 60.0 | + | + | + | + | + | + | + | + |
| *A2BP1: exon 12 (mainly utr) + downstream* | | | | | R: | CCTCCCATAATATAAAAGTCAGCAA | | | | | | | | | |
| GSPT1 | 14: | 7,543,794 | 745 | | F: | ATTCATTTCATGCCCTGCTC | 60.3 | + | + | m | + | + | + | + | + |
| *GSPT1: intron 8* | | | | | R: | AATTGGCAGCCTGATTGGT | | | | | | | | | |
| C9orf78 | 17: | 6,252,552 | 657 | | F: | TGGCAGAGCAGCAGAACA | 60.3 | + | m | + | + | + | + | + | + |
| *C9orf78: intron 7* | | | | | R: | TCTCTGTRTCCCCCACTCTC | | | | | | | | | |
| OLFM1_ds | 17: | 8,674,349 | 778 | | F: | ATTAAGACCAACRTCATGGAGG | 60.2 | + | + | + | + | + | + | + | + |
| *OLFM1_ds: downstream (exon, utr?)* | | | | | R: | GATTTCCAACCACCATAATGCTA | | | | | | | | | |
| SPAG9 | 18: | 9,602,824 | 319 | 8 | F: | AYGCACAGCTYTGCTTCC | 60.0 | + | + | + | + | + | + | + | + |
| *SPAG9: intron 35* | | | | | R: | CCRCTTATCACCAACATTGAT | | | | | | | | | |

| | | | | | | | | | | | | | | | |
|---|---|---|---|---|---|---|---|---|---|---|---|---|---|---|---|
| ENSGALT00000001658 | 19: | 770,102 | 627 | 1 | F: | ATTGAACGTGCTCACATGAG | 63–53 | + | + | + | – | + | + | + | + |
| *SBDS: intron 2* | | | | | R: | TTCCTCCACATCTTTCAGAC | | | | | | | | | |
| ENSGALT00000005087 | 19: | 6,981,967 | 519 | 1 | F: | CTGGCAAATCATGATGACAG | 63–53 | + | + | + | + | + | + | + | + |
| *POLDIP2: intron 5* | | | | | R: | CTGGATGGGCACTTGATCAG | | | | | | | | | |
| βact3 | 22: | 735,035 | 334 | 6 | F: | GGCAATGAGAGGTTCAGGT | 62.0 | + | + | + | + | + | + | + | + |
| *Q90WZ7_TAEGU: intron 4* | | | | | R: | TGGTACCACCAGACAGCAC | | | | | | | | | |
| α-B crystallin | 24: | 1,639,781 | 117 | 2 | F: | CTGATCCGCAGACCTTTCTT | 59.0 | + | + | + | + | + | + | + | –/m |
| *CRYAB: exon 1* | | | | | R: | TAGCCAACTGGGCATCCGC | | | | | | | | | |
| CEPUS | 24: | 6,314,469 | 810 | 2 | F: | CGAGTCAAAGTCACCGTCAA | 64.0 | + | m | + | + | + | + | + | + |
| *NTM: intron 4* | | | | | R: | CTCTTCGCATCCGAGATGTA | | | | | | | | | |
| ACL | 27: | 2,282,159 | 463 | 2 | F: | GCTCTGCTTATGACAGCACT | 56.0 | + | + | + | m | + | + | + | + |
| *ACL: intron 18* | | | | | R: | CAGCAATAATGGCAATGGTG | | | | | | | | | |
| ENSGALT00000001183 | 28: | 1,722,106 | 519 | 1 | F: | CAGTCAAGGCTTACAGATGC | 63–53 | + | + | + | + | + | + | + | + |
| *OAZ1: intron 3* | | | | | R: | ATCTCTGTTCTTGTGGAAAC | | | | | | | | | |
| SEMA6A_ds | Z_random: | 2,321,651 | 732 | | F: | GCTAACTCTGACTTAACAATGGCA | 59.8 | + | + | m | + | + | + | + | + |
| *SEMA6A: downstream (exon, utr?)* | | | | | R: | AGGGTGTAAGGCACAAATTTACA | | | | | | | | | |
| LAMA | Un: | 106,966,369 | 89 | 2, 7 | F: | CCAAGAAGCAGCTGCAGGATGAGATGC | 69.0 | + | + | + | + | + | + | + | + |
| *LMNA-2: intron 2* | | | | | R: | CTGCCGCCCGTTGTCGATCTCCACCAG | | | | | | | | | |

*References: 1 – Backström *et al.* (2008); 2 – Primmer *et al.* (2002); 3 – Heslewood *et al.* (1998); 4 – Slade *et al.* (1993); 5 – Friesen *et al.* (1999); 6 – Carling & Brumfield (2009); 7 – Friesen *et al.* (1997); 8 – Stervander *et al.* (2015).

†Species included in test panel: great reed warbler *Acrocephalus arundinaceus*, blue tit *Cyanistes caeruleus*, hooded crow *Corvus corone cornix*, zebra finch *Taeniopygia guttata*, yellow canary *Crithagra flaviventris*, thick-billed seedeater *Crithagra burtoni*, Príncipe seedeater *Crithagra rufobrunnea*, and Inaccessible Island finch *Nesospiza acunhae*.

[a] ENSGALT00000025969 should also amplify the Tgu paralogue ENSTGUG00000015403 (Tgu Un: 155,130,434-155,134,099).
[b] ANKRD17 amplify the complimentary strand to the gene.
[c] The product of ENSGALT00000011836 corresponds to a segment which is duplicated within the Tgu PSMD6 intron 5.

Table S8. Matrix characteristics and substitution models for the nuclear sequence markers. Number of samples are presented for the ingroup.

| | Matrix length (bp) | | | | |
|---|---|---|---|---|---|
| **Marker** | ingroup | total | **N samples** | **Substitution model** | **Across-site mutation rates**[a] |
| AKIRIN2 | 709 | 709 | 32 | GTR | equal |
| ALG11 | 682 | 721 | 32 | GTR | equal |
| BTF3L4 | 355 | 366 | 32 | HKY85 | equal |
| C9orf78 | 333 | 334 | 31 | HKY85 | equal |
| CEPUS | 771 | 775 | 32 | HKY85 | equal |
| CHP | 440 | 441 | 30 | HKY85 | gamma distr. |
| CNN3_ds | 675 | 676 | 32 | GTR | equal |
| DPP10_ds | 595 | 605 | 32 | HKY85 | equal |
| EIF4G2 | 777 | 782 | 32 | HKY85 | prop. inv. |
| ENSGALT00000001183 | 485 | 537 | 30 | HKY85 | prop. inv. |
| ENSGALT00000001658 | 641 | 654 | 31 | GTR | equal |
| ENSGALT00000005087 | 576 | 578 | 32 | HKY85 | equal |
| ENSGALT00000015724 | 673 | 695 | 32 | HKY85 | prop. inv. |
| ENSGALT00000016264 | 654 | 689 | 32 | HKY85 | prop. inv. |
| ENSGALT00000018851 | 731 | 753 | 32 | HKY85 | equal |
| ENSGALT00000020352 | 501 | 608 | 32 | HKY85 | equal |
| ENSGALT00000026187 | 778 | 818 | 29 | HKY85 | prop. inv. |
| ENSGALT00000026472 | 929 | 946 | 32 | HKY85 | equal |
| ENSGALT00000027331 | 627 | 629 | 32 | GTR | equal |
| GADPH | 230 | 238 | 30 | GTR | equal |
| GSPT1 | 732 | 751 | 32 | HKY85 | prop. inv. |
| LRPP4/RP40 | 244 | 246 | 32 | K80 | equal |
| Myo2 | 659 | 671 | 32 | K80 | equal |
| OGDHL | 702 | 714 | 32 | HKY85 | equal |
| PCDH18 | 950 | 955 | 32 | GTR | equal |
| PPP2CA | 506 | 506 | 32 | F81 | equal |
| Rho1 | 903 | 918 | 32 | F81 | equal |
| RNF146 | 461 | 461 | 32 | GTR | equal |
| RPL7 | 638 | 648 | 31 | HKY85 | prop. inv. |
| SNAP25 | 795 | 800 | 32 | HKY85 | prop. inv. |
| Tgu1A-49.14MbpEx | 319 | 321 | 32 | F81 | equal |
| WDR33 | 627 | 646 | 32 | HKY85 | equal |
| WDR47 | 296 | 315 | 32 | HKY85 | equal |

[a] Across-site mutation rates are either set as equal, a proportion of the sites being invariable and the remaining sites at equal rates (prop. inv.), or variable, where mutation rates for a site is drawn from a gamma distribution.

**Table S9.** Data accession for mitochondrial DNA (mtDNA) and nuclear DNA (nuDNA) sequences (GenBank accession numbers), and restriction site-associated DNA (RAD) sequencing single nucleotide polymorphism (SNP) data (Sequence Read Archive run id). As identifiers, the voucher (FIELD in Table S5) and the names used in different trees are listed. The table is also available as a spreadsheet in the Zenodo data deposition (https://doi.org/10.5281/zenodo.5816808).

[Table starts on next page.]

| Sample | Sample name in tree/dataset nuDNA | mtDNA | RAD | RADseq | Mt Haplotypes | Mt markers |
|---|---|---|---|---|---|---|
| *C. r. rufobrunnea* | | | | | | |
| P1_141 | SRR4 | P001 | | | Ruf3 | KY078928 |
| P1_142 | SRR8 | P002 | | | Ruf4 | KY078929 |
| P1_266 | | P003 | | | Ruf5 | OL441099 |
| P1_267 | SRR7 | P004 | | | Ruf5 | OL441098 |
| P2_021 | SRR5 | P005 | | | Ruf3 | OL441095 |
| P3_026 | | P011 | | | Ruf5 | OL441097 |
| P3_027 | SRR6 | P012 | | | Ruf3 | OL441096 |
| P3_030 | | P013 | | | Ruf6 | OL441092 |
| P3_031 | SRR10 | P014 | | | Ruf3 | OL441094 |
| P3_032 | | P015 | | | Ruf3 | OL441093 |
| P3_041 | SRR9 | | | | | |
| P3_045 | SRR2 | | | | | |
| P3_055 | SRR3 | | | | | |
| P3_057 | SRR1 | | | | | |
| P5_030 | | | P1 | SRR17333565 | | |
| P5_031 | | | P2 | SRR17333564 | | |
| P5_034 | | | P3 | SRR17333553 | | |
| P5_035 | | | P4 | SRR17333542 | | |
| P5_038 | | | P5 | SRR17333541 | | |
| P5_042 | | | P6 | SRR17333540 | | |
| P5_045 | | | P7 | SRR17333539 | | |
| P3_058/P5_047 | | | P8 | SRR17333538 | | |
| *C. r. fradei* | | | | | | |
| P1_230 | | B001 | | | Ruf2 | KY078930 |
| P1_231 | | B002 | | | Ruf1 | KY078931 |
| P1_232 | | B003 | | | Ruf1 | OL441104 |
| P1_233 | | B004 | | | Ruf2 | OL441107 |
| P1_234 | SRF7 | B005 | | | Ruf2 | OL441108 |
| P1_235 | | B006 | | | Ruf2 | OL441106 |
| P1_236 | SRF6 | B007 | | | Ruf1 | OL441103 |
| P1_239 | SRF5 | B010 | | | Ruf1 | OL441102 |
| P1_243 | | B014 | | | Ruf1 | OL441101 |
| P1_249 | | B019 | | | Ruf2 | OL441105 |
| P1_250 | SRF4 | B020 | | | Ruf1 | OL441100 |
| P1_259 | SRF1 | | | | | |
| P1_263 | SRF2 | | | | | |
| P2_001 | SRF9 | | | | | |
| P3_001 | SRF8 | | | | | |
| P3_004 | SRF3 | | | | | |
| P5_005 | | | B5 | SRR17333537 | | |
| P5_006 | | | B6 | SRR17333536 | | |
| P5_011 | | | B7 | SRR17333563 | | |
| P5_016 | | | B8 | SRR17333562 | | |
| P5_010 | | | B1 | SRR17333561 | | |
| P5_014 | | | B2 | SRR17333560 | | |
| P5_015 | | | B3 | SRR17333559 | | |
| P5_028 | | | B4 | SRR17333558 | | |

| Sample | Sample name in tree/dataset nuDNA | mtDNA | RAD | RADseq | Mt Haplotypes | Mt markers |
|---|---|---|---|---|---|---|
| *C. r. thomensis* | | | | | | |
| ST1_044 | SRT3 | ST005 | | | Ruf7 | OL441117 |
| ST1_100 | SRT2 | ST008 | | | Ruf8 | OL441113 |
| ST1_125/ST5_027 | | ST015 | ST1 | SRR17333557 | Ruf9 | OL441109 |
| ST1_146 | SRT1 | ST016 | | | Ruf4 | KY078933 |
| ST1_204 | SRT5 | ST017 | | | Ruf10 | OL441115 |
| ST1_225 | | ST020 | | | Ruf9 | KY078932 |
| ST1_242 | | ST024 | | | Ruf11 | OL441112 |
| ST1_312 | SRT6 | | | | | |
| ST2_108 | SRT9 | | | | | |
| ST2_157 | SRT7 | ST068 | | | Ruf7 | OL441116 |
| ST2_256 | SRT10 | ST072 | | | Ruf12 | OL441110 |
| ST3_012 | SRT4 | | | | | |
| ST3_015 | | ST077 | | | Ruf4 | OL441114 |
| ST3_295 | | ST112 | | | Ruf13 | OL441111 |
| ST3_342 | SRT8 | | | | | |
| ST5_023 | | | ST2 | SRR17333556 | | |
| ST5_026 | | | ST8 | SRR17333555 | | |
| ST5_097 | | | ST3 | SRR17333554 | | |
| ST5_098 | | | ST4 | SRR17333552 | | |
| ST5_100 | | | ST5 | SRR17333551 | | |
| ST5_123 | | | ST6 | SRR17333550 | | |
| ST5_125 | | | ST7 | SRR17333549 | | |
| *C. concolor* | | | | | | |
| ST1_362 | NC1 | Neo1 | Neo3 | | Neo1 | KY078934 |
| ST3_318 | NC2 | Neo2 | Neo4 | | Neo2 | KY078935 |
| ST3_319 | NC3 | Neo3 | Neo2 | | Neo3 | KY078936 |
| ST5_039 | NC4 | | Neo1 | | | |
| *C. burtoni* | | | | | | |
| C084 | | Bur1 | | | | KY078926 |
| C119 | | Bur2 | | | | KY078927 |
| C133 | SB2 | | | | | |
| C159 | SB1 | | | | | |
| *C. flaviventris* | | | | | | |
| Z108 | SF1 | | | | | |
| Z111 | SF2 | | | | | |

| | Nucelar markers | | | | | | |
|---|---|---|---|---|---|---|---|
| **Sample** | ACADSB | ACTR6 | AKIRIN2 | ALG11 | ATP6AP2 | BTF3L4 | C9orf78 |
| *C. r. rufobrunnea* | | | | | | | |
| P1_141 | MG643135 | MG643170 | MG643206 | MG643241 | MG643274 | MG643309 | MG643344 |
| P1_142 | MG643139 | MG643174 | MG643210 | MG643245 | MG643277 | MG643313 | MG643348 |
| P1_266 | | | | | | | |
| P1_267 | MG643138 | MG643173 | MG643209 | MG643244 | – | MG643312 | MG643347 |
| P2_021 | MG643136 | MG643171 | MG643207 | MG643242 | MG643275 | MG643310 | MG643345 |
| P3_026 | | | | | | | |
| P3_027 | MG643137 | MG643172 | MG643208 | MG643243 | MG643276 | MG643311 | MG643346 |
| P3_030 | | | | | | | |
| P3_031 | MG643141 | MG643176 | MG643212 | MG643247 | MG643279 | MG643315 | MG643350 |
| P3_032 | | | | | | | |
| P3_041 | MG643140 | MG643175 | MG643211 | MG643246 | MG643278 | MG643314 | MG643349 |
| P3_045 | MG643133 | MG643168 | MG643204 | MG643239 | MG643272 | MG643307 | MG643342 |
| P3_055 | MG643134 | MG643169 | MG643205 | MG643240 | MG643273 | MG643308 | MG643343 |
| P3_057 | MG643132 | MG643167 | MG643203 | MG643238 | MG643271 | MG643306 | MG643341 |
| P5_030 | | | | | | | |
| P5_031 | | | | | | | |
| P5_034 | | | | | | | |
| P5_035 | | | | | | | |
| P5_038 | | | | | | | |
| P5_042 | | | | | | | |
| P5_045 | | | | | | | |
| P3_058/P5_047 | | | | | | | |
| *C. r. fradei* | | | | | | | |
| P1_230 | | | | | | | |
| P1_231 | | | | | | | |
| P1_232 | | | | | | | |
| P1_233 | | | | | | | |
| P1_234 | MG643129 | MG643164 | MG643200 | MG643235 | MG643268 | MG643303 | MG643338 |
| P1_235 | | | | | | | |
| P1_236 | MG643128 | MG643163 | MG643199 | MG643234 | MG643267 | MG643302 | MG643337 |
| P1_239 | MG643127 | MG643162 | MG643198 | MG643233 | – | MG643301 | MG643336 |
| P1_243 | | | | | | | |
| P1_249 | | | | | | | |
| P1_250 | MG643126 | MG643161 | MG643197 | MG643232 | MG643266 | MG643300 | MG643335 |
| P1_259 | MG643123 | MG643158 | MG643194 | MG643229 | MG643264 | MG643297 | MG643332 |
| P1_263 | MG643124 | MG643159 | MG643195 | MG643230 | MG643265 | MG643298 | MG643333 |
| P2_001 | MG643131 | MG643166 | MG643202 | MG643237 | MG643270 | MG643305 | MG643340 |
| P3_001 | MG643130 | MG643165 | MG643201 | MG643236 | MG643269 | MG643304 | MG643339 |
| P3_004 | MG643125 | MG643160 | MG643196 | MG643231 | – | MG643299 | MG643334 |
| P5_005 | | | | | | | |
| P5_006 | | | | | | | |
| P5_011 | | | | | | | |
| P5_016 | | | | | | | |
| P5_010 | | | | | | | |
| P5_014 | | | | | | | |
| P5_015 | | | | | | | |
| P5_028 | | | | | | | |

| | Nucelar markers | | | | | | |
|---|---|---|---|---|---|---|---|
| **Sample** | ACADSB | ACTR6 | AKIRIN2 | ALG11 | ATP6AP2 | BTF3L4 | C9orf78 |
| *C. r. thomensis* | | | | | | | |
| ST1_044 | MG643144 | MG643179 | MG643215 | MG643250 | MG643282 | MG643318 | MG643352 |
| ST1_100 | MG643143 | MG643178 | MG643214 | MG643249 | MG643281 | MG643317 | MG643351 |
| ST1_125/ST5_027 | | | | | | | |
| ST1_146 | MG643142 | MG643177 | MG643213 | MG643248 | MG643280 | MG643316 | – |
| ST1_204 | MG643146 | MG643181 | MG643217 | MG643252 | MG643284 | MG643320 | MG643354 |
| ST1_225 | | | | | | | |
| ST1_242 | | | | | | | |
| ST1_312 | MG643147 | MG643182 | MG643218 | MG643253 | MG643285 | MG643321 | MG643355 |
| ST2_108 | MG643150 | MG643185 | MG643221 | MG643256 | MG643288 | MG643324 | MG643358 |
| ST2_157 | MG643148 | MG643183 | MG643219 | MG643254 | MG643286 | MG643322 | MG643356 |
| ST2_256 | MG643151 | MG643186 | MG643222 | MG643257 | MG643289 | MG643325 | MG643359 |
| ST3_012 | MG643145 | MG643180 | MG643216 | MG643251 | MG643283 | MG643319 | MG643353 |
| ST3_015 | | | | | | | |
| ST3_295 | | | | | | | |
| ST3_342 | MG643149 | MG643184 | MG643220 | MG643255 | MG643287 | MG643323 | MG643357 |
| ST5_023 | | | | | | | |
| ST5_026 | | | | | | | |
| ST5_097 | | | | | | | |
| ST5_098 | | | | | | | |
| ST5_100 | | | | | | | |
| ST5_123 | | | | | | | |
| ST5_125 | | | | | | | |
| *C. concolor* | | | | | | | |
| ST1_362 | | | | | | | |
| ST3_318 | | | | | | | |
| ST3_319 | | | | | | | |
| ST5_039 | | | | | | | |
| *C. burtoni* | | | | | | | |
| C084 | | | | | | | |
| C119 | | | | | | | |
| C133 | MG643120 | MG643156 | MG643191 | MG643227 | MG643262 | MG643294 | MG643330 |
| C159 | MG643119 | MG643155 | MG643190 | MG643226 | MG643261 | MG643293 | MG643329 |
| *C. flaviventris* | | | | | | | |
| Z108 | MG643121 | MG643157 | MG643192 | MG643228 | – | MG643295 | – |
| Z111 | MG643122 | – | MG643193 | – | MG643263 | MG643296 | MG643331 |

| Sample | Nucelar markers | | | | | | |
|---|---|---|---|---|---|---|---|
| | CHP1 | CNN3 | COL14A1 | CRIPT | DPP10 | EIF4G2 | GAPDH |
| *C. r. rufobrunnea* | | | | | | | |
| P1_141 | MG643379 | MG643413 | MG643449 | MG643484 | MG643520 | MG643556 | MG643590 |
| P1_142 | MG643383 | MG643417 | MG643453 | MG643488 | MG643524 | MG643560 | MG643594 |
| P1_266 | | | | | | | |
| P1_267 | MG643382 | MG643416 | MG643452 | MG643487 | MG643523 | MG643559 | MG643593 |
| P2_021 | MG643380 | MG643414 | MG643450 | MG643485 | MG643521 | MG643557 | MG643591 |
| P3_026 | | | | | | | |
| P3_027 | MG643381 | MG643415 | MG643451 | MG643486 | MG643522 | MG643558 | MG643592 |
| P3_030 | | | | | | | |
| P3_031 | MG643385 | MG643419 | MG643455 | MG643490 | MG643526 | MG643562 | MG643596 |
| P3_032 | | | | | | | |
| P3_041 | MG643384 | MG643418 | MG643454 | MG643489 | MG643525 | MG643561 | MG643595 |
| P3_045 | MG643377 | MG643411 | MG643447 | MG643482 | MG643518 | MG643554 | – |
| P3_055 | MG643378 | MG643412 | MG643448 | MG643483 | MG643519 | MG643555 | MG643589 |
| P3_057 | MG643376 | MG643410 | MG643446 | MG643481 | MG643517 | MG643553 | MG643588 |
| P5_030 | | | | | | | |
| P5_031 | | | | | | | |
| P5_034 | | | | | | | |
| P5_035 | | | | | | | |
| P5_038 | | | | | | | |
| P5_042 | | | | | | | |
| P5_045 | | | | | | | |
| P3_058/P5_047 | | | | | | | |
| *C. r. fradei* | | | | | | | |
| P1_230 | | | | | | | |
| P1_231 | | | | | | | |
| P1_232 | | | | | | | |
| P1_233 | | | | | | | |
| P1_234 | MG643373 | MG643407 | MG643443 | MG643478 | MG643514 | MG643550 | MG643585 |
| P1_235 | | | | | | | |
| P1_236 | MG643372 | MG643406 | MG643442 | MG643477 | MG643513 | MG643549 | MG643584 |
| P1_239 | MG643371 | MG643405 | MG643441 | MG643476 | MG643512 | MG643548 | MG643583 |
| P1_243 | | | | | | | |
| P1_249 | | | | | | | |
| P1_250 | MG643370 | MG643404 | MG643440 | MG643475 | MG643511 | MG643547 | MG643582 |
| P1_259 | MG643367 | MG643401 | MG643437 | MG643472 | MG643508 | MG643544 | MG643580 |
| P1_263 | MG643368 | MG643402 | MG643438 | MG643473 | MG643509 | MG643545 | – |
| P2_001 | MG643375 | MG643409 | MG643445 | MG643480 | MG643516 | MG643552 | MG643587 |
| P3_001 | MG643374 | MG643408 | MG643444 | MG643479 | MG643515 | MG643551 | MG643586 |
| P3_004 | MG643369 | MG643403 | MG643439 | MG643474 | MG643510 | MG643546 | MG643581 |
| P5_005 | | | | | | | |
| P5_006 | | | | | | | |
| P5_011 | | | | | | | |
| P5_016 | | | | | | | |
| P5_010 | | | | | | | |
| P5_014 | | | | | | | |
| P5_015 | | | | | | | |
| P5_028 | | | | | | | |

| Sample | Nucelar markers | | | | | | |
|---|---|---|---|---|---|---|---|
| | CHP1 | CNN3 | COL14A1 | CRIPT | DPP10 | EIF4G2 | GAPDH |
| *C. r. thomensis* | | | | | | | |
| ST1_044 | – | MG643422 | MG643458 | MG643493 | MG643529 | MG643565 | MG643599 |
| ST1_100 | – | MG643421 | MG643457 | MG643492 | MG643528 | MG643564 | MG643598 |
| ST1_125/ST5_027 | | | | | | | |
| ST1_146 | MG643386 | MG643420 | MG643456 | MG643491 | MG643527 | MG643563 | MG643597 |
| ST1_204 | MG643388 | MG643424 | MG643460 | MG643495 | MG643531 | MG643567 | MG643601 |
| ST1_225 | | | | | | | |
| ST1_242 | | | | | | | |
| ST1_312 | MG643389 | MG643425 | MG643461 | MG643496 | MG643532 | MG643568 | MG643602 |
| ST2_108 | MG643392 | MG643428 | MG643464 | MG643499 | MG643535 | MG643571 | MG643605 |
| ST2_157 | MG643390 | MG643426 | MG643462 | MG643497 | MG643533 | MG643569 | MG643603 |
| ST2_256 | MG643393 | MG643429 | MG643465 | MG643500 | MG643536 | MG643572 | MG643606 |
| ST3_012 | MG643387 | MG643423 | MG643459 | MG643494 | MG643530 | MG643566 | MG643600 |
| ST3_015 | | | | | | | |
| ST3_295 | | | | | | | |
| ST3_342 | MG643391 | MG643427 | MG643463 | MG643498 | MG643534 | MG643570 | MG643604 |
| ST5_023 | | | | | | | |
| ST5_026 | | | | | | | |
| ST5_097 | | | | | | | |
| ST5_098 | | | | | | | |
| ST5_100 | | | | | | | |
| ST5_123 | | | | | | | |
| ST5_125 | | | | | | | |
| *C. concolor* | | | | | | | |
| ST1_362 | | | | | | | |
| ST3_318 | | | | | | | |
| ST3_319 | | | | | | | |
| ST5_039 | | | | | | | |
| *C. burtoni* | | | | | | | |
| C084 | | | | | | | |
| C119 | | | | | | | |
| C133 | MG643364 | MG643398 | MG643434 | MG643470 | MG643505 | MG643541 | MG643577 |
| C159 | MG643363 | MG643397 | MG643433 | MG643469 | MG643504 | MG643540 | MG643576 |
| *C. flaviventris* | | | | | | | |
| Z108 | MG643365 | MG643399 | MG643435 | MG643471 | MG643506 | MG643542 | MG643578 |
| Z111 | MG643366 | MG643400 | MG643436 | – | MG643507 | MG643543 | MG643579 |

| | Nucelar markers | | | | | | |
|---|---|---|---|---|---|---|---|
| **Sample** | GSPT1 | HNT | HUS1 | MB | OAZ1 | OGDHL | PCDH18 |
| *C. r. rufobrunnea* | | | | | | | |
| P1_141 | MG643626 | MG643765 | MG643661 | MG643730 | MG643799 | MG643835 | MG643870 |
| P1_142 | MG643630 | MG643769 | MG643665 | MG643734 | MG643803 | MG643839 | MG643874 |
| P1_266 | | | | | | | |
| P1_267 | MG643629 | MG643768 | MG643664 | MG643733 | MG643802 | MG643838 | MG643873 |
| P2_021 | MG643627 | MG643766 | MG643662 | MG643731 | MG643800 | MG643836 | MG643871 |
| P3_026 | | | | | | | |
| P3_027 | MG643628 | MG643767 | MG643663 | MG643732 | MG643801 | MG643837 | MG643872 |
| P3_030 | | | | | | | |
| P3_031 | MG643632 | MG643771 | MG643667 | MG643736 | MG643805 | MG643841 | MG643876 |
| P3_032 | | | | | | | |
| P3_041 | MG643631 | MG643770 | MG643666 | MG643735 | MG643804 | MG643840 | MG643875 |
| P3_045 | MG643624 | MG643763 | MG643659 | MG643728 | MG643797 | MG643833 | MG643868 |
| P3_055 | MG643625 | MG643764 | MG643660 | MG643729 | MG643798 | MG643834 | MG643869 |
| P3_057 | MG643623 | MG643762 | MG643658 | MG643727 | MG643796 | MG643832 | MG643867 |
| P5_030 | | | | | | | |
| P5_031 | | | | | | | |
| P5_034 | | | | | | | |
| P5_035 | | | | | | | |
| P5_038 | | | | | | | |
| P5_042 | | | | | | | |
| P5_045 | | | | | | | |
| P3_058/P5_047 | | | | | | | |
| *C. r. fradei* | | | | | | | |
| P1_230 | | | | | | | |
| P1_231 | | | | | | | |
| P1_232 | | | | | | | |
| P1_233 | | | | | | | |
| P1_234 | MG643620 | MG643759 | MG643655 | MG643724 | – | MG643829 | MG643864 |
| P1_235 | | | | | | | |
| P1_236 | MG643619 | MG643758 | MG643654 | MG643723 | MG643793 | MG643828 | MG643863 |
| P1_239 | MG643618 | MG643757 | MG643653 | MG643722 | MG643792 | MG643827 | MG643862 |
| P1_243 | | | | | | | |
| P1_249 | | | | | | | |
| P1_250 | MG643617 | MG643756 | MG643652 | MG643721 | – | MG643826 | MG643861 |
| P1_259 | MG643614 | MG643753 | MG643649 | MG643718 | MG643789 | MG643823 | MG643858 |
| P1_263 | MG643615 | MG643754 | MG643650 | MG643719 | MG643790 | MG643824 | MG643859 |
| P2_001 | MG643622 | MG643761 | MG643657 | MG643726 | MG643795 | MG643831 | MG643866 |
| P3_001 | MG643621 | MG643760 | MG643656 | MG643725 | MG643794 | MG643830 | MG643865 |
| P3_004 | MG643616 | MG643755 | MG643651 | MG643720 | MG643791 | MG643825 | MG643860 |
| P5_005 | | | | | | | |
| P5_006 | | | | | | | |
| P5_011 | | | | | | | |
| P5_016 | | | | | | | |
| P5_010 | | | | | | | |
| P5_014 | | | | | | | |
| P5_015 | | | | | | | |
| P5_028 | | | | | | | |

| Sample | Nucelar markers | | | | | | |
|---|---|---|---|---|---|---|---|
| | GSPT1 | HNT | HUS1 | MB | OAZ1 | OGDHL | PCDH18 |
| *C. r. thomensis* | | | | | | | |
| ST1_044 | MG643635 | MG643774 | MG643670 | MG643739 | MG643808 | MG643844 | MG643879 |
| ST1_100 | MG643634 | MG643773 | MG643669 | MG643738 | MG643807 | MG643843 | MG643878 |
| ST1_125/ST5_027 | | | | | | | |
| ST1_146 | MG643633 | MG643772 | MG643668 | MG643737 | MG643806 | MG643842 | MG643877 |
| ST1_204 | MG643637 | MG643776 | MG643672 | MG643741 | MG643810 | MG643846 | MG643881 |
| ST1_225 | | | | | | | |
| ST1_242 | | | | | | | |
| ST1_312 | MG643638 | MG643777 | MG643673 | MG643742 | MG643811 | MG643847 | MG643882 |
| ST2_108 | MG643641 | MG643780 | MG643676 | MG643745 | MG643814 | MG643850 | MG643885 |
| ST2_157 | MG643639 | MG643778 | MG643674 | MG643743 | MG643812 | MG643848 | MG643883 |
| ST2_256 | MG643642 | MG643781 | MG643677 | MG643746 | MG643815 | MG643851 | MG643886 |
| ST3_012 | MG643636 | MG643775 | MG643671 | MG643740 | MG643809 | MG643845 | MG643880 |
| ST3_015 | | | | | | | |
| ST3_295 | | | | | | | |
| ST3_342 | MG643640 | MG643779 | MG643675 | MG643744 | MG643813 | MG643849 | MG643884 |
| ST5_023 | | | | | | | |
| ST5_026 | | | | | | | |
| ST5_097 | | | | | | | |
| ST5_098 | | | | | | | |
| ST5_100 | | | | | | | |
| ST5_123 | | | | | | | |
| ST5_125 | | | | | | | |
| *C. concolor* | | | | | | | |
| ST1_362 | | | | | | | |
| ST3_318 | | | | | | | |
| ST3_319 | | | | | | | |
| ST5_039 | | | | | | | |
| *C. burtoni* | | | | | | | |
| C084 | | | | | | | |
| C119 | | | | | | | |
| C133 | MG643611 | MG643751 | MG643646 | – | MG643786 | MG643820 | MG643856 |
| C159 | MG643610 | MG643750 | – | – | MG643785 | MG643819 | MG643855 |
| *C. flaviventris* | | | | | | | |
| Z108 | MG643612 | – | MG643647 | MG643716 | MG643787 | MG643821 | – |
| Z111 | MG643613 | MG643752 | MG643648 | MG643717 | MG643788 | MG643822 | MG643857 |

| Sample | Nucelar markers | | | | | | |
|---|---|---|---|---|---|---|---|
| | POLDIP2 | PPP2CA | RBM26 | RHO-2 | RNF146 | RPL7 | RPSA |
| *C. r. rufobrunnea* | | | | | | | |
| P1_141 | MG643906 | MG643942 | MG643977 | MG644011 | MG644047 | MG644083 | MG644118 |
| P1_142 | MG643910 | MG643946 | MG643981 | MG644015 | MG644051 | MG644087 | MG644122 |
| P1_266 | | | | | | | |
| P1_267 | MG643909 | MG643945 | MG643980 | MG644014 | MG644050 | MG644086 | MG644121 |
| P2_021 | MG643907 | MG643943 | MG643978 | MG644012 | MG644048 | MG644084 | MG644119 |
| P3_026 | | | | | | | |
| P3_027 | MG643908 | MG643944 | MG643979 | MG644013 | MG644049 | MG644085 | MG644120 |
| P3_030 | | | | | | | |
| P3_031 | MG643912 | MG643948 | MG643983 | MG644017 | MG644053 | MG644089 | MG644124 |
| P3_032 | | | | | | | |
| P3_041 | MG643911 | MG643947 | MG643982 | MG644016 | MG644052 | MG644088 | MG644123 |
| P3_045 | MG643904 | MG643940 | MG643975 | MG644009 | MG644045 | MG644081 | MG644116 |
| P3_055 | MG643905 | MG643941 | MG643976 | MG644010 | MG644046 | MG644082 | MG644117 |
| P3_057 | MG643903 | MG643939 | MG643974 | MG644008 | MG644044 | MG644080 | MG644115 |
| P5_030 | | | | | | | |
| P5_031 | | | | | | | |
| P5_034 | | | | | | | |
| P5_035 | | | | | | | |
| P5_038 | | | | | | | |
| P5_042 | | | | | | | |
| P5_045 | | | | | | | |
| P3_058/P5_047 | | | | | | | |
| *C. r. fradei* | | | | | | | |
| P1_230 | | | | | | | |
| P1_231 | | | | | | | |
| P1_232 | | | | | | | |
| P1_233 | | | | | | | |
| P1_234 | MG643900 | MG643936 | MG643971 | MG644005 | MG644041 | MG644077 | MG644112 |
| P1_235 | | | | | | | |
| P1_236 | MG643899 | MG643935 | MG643970 | MG644004 | MG644040 | MG644076 | MG644111 |
| P1_239 | MG643898 | MG643934 | MG643969 | MG644003 | MG644039 | MG644075 | MG644110 |
| P1_243 | | | | | | | |
| P1_249 | | | | | | | |
| P1_250 | MG643897 | MG643933 | MG643968 | MG644002 | MG644038 | MG644074 | MG644109 |
| P1_259 | MG643894 | MG643930 | MG643965 | MG643999 | MG644035 | MG644071 | MG644106 |
| P1_263 | MG643895 | MG643931 | MG643966 | MG644000 | MG644036 | MG644072 | MG644107 |
| P2_001 | MG643902 | MG643938 | MG643973 | MG644007 | MG644043 | MG644079 | MG644114 |
| P3_001 | MG643901 | MG643937 | MG643972 | MG644006 | MG644042 | MG644078 | MG644113 |
| P3_004 | MG643896 | MG643932 | MG643967 | MG644001 | MG644037 | MG644073 | MG644108 |
| P5_005 | | | | | | | |
| P5_006 | | | | | | | |
| P5_011 | | | | | | | |
| P5_016 | | | | | | | |
| P5_010 | | | | | | | |
| P5_014 | | | | | | | |
| P5_015 | | | | | | | |
| P5_028 | | | | | | | |

| | Nucelar markers | | | | | | |
|---|---|---|---|---|---|---|---|
| **Sample** | POLDIP2 | PPP2CA | RBM26 | RHO-2 | RNF146 | RPL7 | RPSA |
| *C. r. thomensis* | | | | | | | |
| ST1_044 | MG643915 | MG643951 | MG643986 | MG644020 | MG644056 | MG644091 | MG644127 |
| ST1_100 | MG643914 | MG643950 | MG643985 | MG644019 | MG644055 | MG644090 | MG644126 |
| ST1_125/ST5_027 | | | | | | | |
| ST1_146 | MG643913 | MG643949 | MG643984 | MG644018 | MG644054 | – | MG644125 |
| ST1_204 | MG643917 | MG643953 | MG643988 | MG644022 | MG644058 | MG644093 | MG644129 |
| ST1_225 | | | | | | | |
| ST1_242 | | | | | | | |
| ST1_312 | MG643918 | MG643954 | MG643989 | MG644023 | MG644059 | MG644094 | MG644130 |
| ST2_108 | MG643921 | MG643957 | MG643992 | MG644026 | MG644062 | MG644097 | MG644133 |
| ST2_157 | MG643919 | MG643955 | MG643990 | MG644024 | MG644060 | MG644095 | MG644131 |
| ST2_256 | MG643922 | MG643958 | MG643993 | MG644027 | MG644063 | MG644098 | MG644134 |
| ST3_012 | MG643916 | MG643952 | MG643987 | MG644021 | MG644057 | MG644092 | MG644128 |
| ST3_015 | | | | | | | |
| ST3_295 | | | | | | | |
| ST3_342 | MG643920 | MG643956 | MG643991 | MG644025 | MG644061 | MG644096 | MG644132 |
| ST5_023 | | | | | | | |
| ST5_026 | | | | | | | |
| ST5_097 | | | | | | | |
| ST5_098 | | | | | | | |
| ST5_100 | | | | | | | |
| ST5_123 | | | | | | | |
| ST5_125 | | | | | | | |
| *C. concolor* | | | | | | | |
| ST1_362 | | | | | | | |
| ST3_318 | | | | | | | |
| ST3_319 | | | | | | | |
| ST5_039 | | | | | | | |
| *C. burtoni* | | | | | | | |
| C084 | | | | | | | |
| C119 | | | | | | | |
| C133 | MG643891 | MG643927 | MG643963 | – | MG644032 | MG644068 | MG644103 |
| C159 | MG643890 | MG643926 | MG643962 | MG643997 | MG644031 | MG644067 | MG644102 |
| *C. flaviventris* | | | | | | | |
| Z108 | MG643892 | MG643928 | – | MG643998 | MG644033 | MG644069 | MG644104 |
| Z111 | MG643893 | MG643929 | MG643964 | – | MG644034 | MG644070 | MG644105 |

| | Nucelar markers | | | | |
|---|---|---|---|---|---|
| **Sample** | SBDS | SNAP25 | Intergenic Tgu1A | WDR33 | WDR47 |
| *C. r. rufobrunnea* | | | | | |
| P1_141 | MG644152 | MG644186 | MG643696 | MG644218 | MG644254 |
| P1_142 | MG644156 | – | MG643700 | MG644222 | MG644257 |
| P1_266 | | | | | |
| P1_267 | MG644155 | – | MG643699 | MG644221 | MG644256 |
| P2_021 | MG644153 | MG644187 | MG643697 | MG644219 | – |
| P3_026 | | | | | |
| P3_027 | MG644154 | MG644188 | MG643698 | MG644220 | MG644255 |
| P3_030 | | | | | |
| P3_031 | MG644158 | MG644190 | MG643702 | MG644224 | MG644259 |
| P3_032 | | | | | |
| P3_041 | MG644157 | MG644189 | MG643701 | MG644223 | MG644258 |
| P3_045 | MG644150 | MG644184 | MG643694 | MG644216 | MG644252 |
| P3_055 | MG644151 | MG644185 | MG643695 | MG644217 | MG644253 |
| P3_057 | MG644149 | MG644183 | MG643693 | MG644215 | MG644251 |
| P5_030 | | | | | |
| P5_031 | | | | | |
| P5_034 | | | | | |
| P5_035 | | | | | |
| P5_038 | | | | | |
| P5_042 | | | | | |
| P5_045 | | | | | |
| P3_058/P5_047 | | | | | |
| *C. r. fradei* | | | | | |
| P1_230 | | | | | |
| P1_231 | | | | | |
| P1_232 | | | | | |
| P1_233 | | | | | |
| P1_234 | MG644146 | – | MG643690 | MG644212 | MG644248 |
| P1_235 | | | | | |
| P1_236 | MG644145 | MG644181 | MG643689 | MG644211 | MG644247 |
| P1_239 | MG644144 | MG644180 | MG643688 | MG644210 | MG644246 |
| P1_243 | | | | | |
| P1_249 | | | | | |
| P1_250 | MG644143 | MG644179 | MG643687 | MG644209 | MG644245 |
| P1_259 | – | MG644176 | MG643684 | MG644206 | MG644242 |
| P1_263 | MG644141 | MG644177 | MG643685 | MG644207 | MG644243 |
| P2_001 | MG644148 | – | MG643692 | MG644214 | MG644250 |
| P3_001 | MG644147 | MG644182 | MG643691 | MG644213 | MG644249 |
| P3_004 | MG644142 | MG644178 | MG643686 | MG644208 | MG644244 |
| P5_005 | | | | | |
| P5_006 | | | | | |
| P5_011 | | | | | |
| P5_016 | | | | | |
| P5_010 | | | | | |
| P5_014 | | | | | |
| P5_015 | | | | | |
| P5_028 | | | | | |

| | Nucelar markers | | | | |
|---|---|---|---|---|---|
| **Sample** | SBDS | SNAP25 | Intergenic Tgu1A | WDR33 | WDR47 |
| *C. r. thomensis* | | | | | |
| ST1_044 | MG644161 | – | MG643705 | MG644227 | MG644262 |
| ST1_100 | MG644160 | – | MG643704 | MG644226 | MG644261 |
| ST1_125/ST5_027 | | | | | |
| ST1_146 | MG644159 | MG644191 | MG643703 | MG644225 | MG644260 |
| ST1_204 | MG644163 | MG644193 | MG643707 | MG644229 | MG644264 |
| ST1_225 | | | | | |
| ST1_242 | | | | | |
| ST1_312 | MG644164 | MG644194 | MG643708 | MG644230 | MG644265 |
| ST2_108 | MG644167 | MG644197 | MG643711 | MG644233 | MG644268 |
| ST2_157 | MG644165 | MG644195 | MG643709 | MG644231 | MG644266 |
| ST2_256 | MG644168 | MG644198 | MG643712 | MG644234 | MG644269 |
| ST3_012 | MG644162 | MG644192 | MG643706 | MG644228 | MG644263 |
| ST3_015 | | | | | |
| ST3_295 | | | | | |
| ST3_342 | MG644166 | MG644196 | MG643710 | MG644232 | MG644267 |
| ST5_023 | | | | | |
| ST5_026 | | | | | |
| ST5_097 | | | | | |
| ST5_098 | | | | | |
| ST5_100 | | | | | |
| ST5_123 | | | | | |
| ST5_125 | | | | | |
| *C. concolor* | | | | | |
| ST1_362 | | | | | |
| ST3_318 | | | | | |
| ST3_319 | | | | | |
| ST5_039 | | | | | |
| *C. burtoni* | | | | | |
| C084 | | | | | |
| C119 | | | | | |
| C133 | MG644139 | MG644173 | MG643682 | MG644203 | MG644239 |
| C159 | MG644138 | MG644172 | MG643681 | MG644202 | MG644238 |
| *C. flaviventris* | | | | | |
| Z108 | – | MG644174 | – | MG644204 | MG644240 |
| Z111 | MG644140 | MG644175 | MG643683 | MG644205 | MG644241 |
